# DOT1L Methyltransferase Activity Preserves SOX2-Enhancer Accessibility And Prevents Activation of Repressed Genes In Murine Stem Cells

**DOI:** 10.1101/2020.02.03.931741

**Authors:** F. Ferrari, L. Arrigoni, H. Franz, L. Butenko, E. Trompouki, T. Vogel, T. Manke

## Abstract

**Background:** During cellular differentiation, the chromatin landscape changes dynamically and contributes to the activation of cell-type specific transcriptional programs. Disruptor of telomeric silencing 1-like (DOT1L) is a histone methyltransferase that mediates mono-, di- and trimethylation of lysine 79 of histone H3 (H3K79me1, 2, 3). Its enzymatic activity is critical for driving cellular differentiation into cardiomyocytes, chondrocytes and neurons, from embryonic or other type of stem cells in physiological settings. Ectopic localization of DOT1L in MLL-rearranged leukemias is causative for leukemogenesis and relapse. Little is known about the causal relevance of DOT1L methyltransferase activity in the global chromatin context and how its enzymatic function affects transcriptional and global chromatin states. Recent reports conducted in leukemia cell models have suggested that deposition of H3K79me2 may be critical to preserve histone H3K27 acetylation (ac) and enhancer activity, and to sustain expression of highly transcribed genes. If and to what extent DOT1L affects chromatin states and enhancer activity during physiological differentiation processes is currently unknown.

**Results:** We measure global changes of seven histone modifications during the differentiation process via high-throughput and quantitative ChIP-seq in an *in-vitro* neuronal differentiation model of mouse embryonic stem cells (mESC). We observe that H3K27ac globally decreases, whereas H3K79me2 globally increases during differentiation, while other modifications remain globally unaltered. Pharmacological inhibition of DOT1L in mESC and mESC-derived neural progenitors results in decreased expression of highly transcribed genes and increased expression of normally repressed genes. Acute DOT1L inhibition primes neural progenitors towards a mature differentiation state. Transcriptional downregulation associates with decreased accessibility of enhancers specifically bound by the master regulator SOX2.

**Conclusions:** *In-vitro* neuronal differentiation couples with a genome-wide accumulation of H3K79me2, never described previously in mammalian cells. Acute inhibition of DOT1L is sufficient to initiate a defined transcriptional program, which biases the transcriptome of neural progenitor cells towards neuronal differentiation. H3K79me2 is not generally causative for maintaining transcriptional levels at a genome-wide scale. In contrast, DOT1L inactivation reduces the chromatin accessibility of enhancers bound by SOX2 *in-vivo*, thereby reducing the expression level of a restricted number of genes. Our work establishes that DOT1L activity gates differentiation of progenitors by allowing SOX2-dependent transcription of stemness programs.

## INTRODUCTION

In eukaryotes, nuclear DNA is wrapped around histones, which constitute the building blocks of chromatin. Histones are subject to a variety of covalent and reversible modifications, mostly affecting lysine, serine and arginine residues (e.g. methylation, acetylation etc.). These post-translational modifications (PTM) are added and removed by specific epigenetic enzymes known as “writers” and “erasers” respectively. The combinatorial presence of these modifications on the chromatin template is thought to add a layer of information, known as the histone code, which builds on top of the genetic code (1).

During differentiation, eukaryotic cells undergo large changes affecting their structural, functional and metabolic profiles. The process is accompanied by major rearrangements of the epigenetic and transcriptional profile, which are driven by the synergistic effects of epigenetic enzymes and transcription factors (2).

Epigenetic and transcriptional changes driving neuronal differentiation have been characterized (3, 4), but few efforts aimed towards a comprehensive description of global histone modification dynamics that affect the chromatin of neural committed cells (5). Previous investigations were limited by the use of semi-quantitative and low-throughput methods (e.g. immunoblotting and imaging). Recent developments in quantitative chromatin immunoprecipitation followed by sequencing (ChIP-seq) have overcome these technical limitations and they now allow to detect genome-wide global changes in histone modifications across conditions in a high-throughput manner (6–8).

Various epigenetic enzymes are important for the orchestration of neuronal differentiation (3, 9). Among these, Disruptor of Telomeric silencing 1 Like (DOT1L) has been recently identified as a critical player in the differentiation process (10–13). DOT1L is a highly conserved histone methyltransferase that catalyzes the mono-, di- and trimethylation of lysine 79 of histone H3 (H3K79me1, 2, 3) (14). Since its first characterization in yeast as a disruptor of telomeric silencing upon gain or loss of function (15, 16), the protein has been recognized to be involved in a variety of biological processes, such as cell cycle control (17), DNA repair (14), gene expression (18), differentiation and reprogramming (19). DOT1L regulates cardiomyocyte differentiation and maturation (20, 21) and chondrocyte differentiation (22), while the modulation of its enzymatic activity was shown to be critical for cellular reprogramming efficiency (10). Within the neural lineage, DOT1L prevents premature cell cycle exit and depletion of the neural progenitor pool and it is necessary for proper neuronal differentiation (13,23,24).

DOT1L plays a prominent role in certain forms of leukemia. Interestingly, some studies in this field identified specific perturbations of the chromatin context that manifest upon blocking of DOT1L and indicate crosstalk between H3K79me2 and histone acetylation. Chen et al. show that *Dot1l* knock-down results in the establishment of repressive chromatin states around MLL target genes. This evidence suggests that the presence of H3K79 methylation may be critical to prevent deacetylation through e.g. activity of SIRT1-complexes (25). Loss of DOT1L activity also results in decreased acetylation and reduced frequency of promoter-enhancer interactions at H3K79me2-marked enhancers (26). Currently, it is not clear whether the molecular perturbations described in leukemia are relevant for the differentiation phenotypes described in other model systems, and whether DOT1L activity targets enhancers in physiological developmental settings.

In this work, we use mouse embryonic stem cells (mESC) and their *in-vitro* derived neural progeny (NPC48h) to systematically characterize the global dynamics of the epigenetic landscape during neuronal differentiation (27). For both mESC and NPC48h, we further investigate whether the competitive inhibition of DOT1L with Pinometostat (EPZ5676) affects the establishment of chromatin states and cell-type specific transcriptional programs.

We show that the global levels of H3K79me2 increases genome-wide during the differentiation process. Our data indicate that DOT1L inactivation causes the onset of a transcriptional program which primes mESC-derived NPC towards neuronal differentiation. We further show that acute DOT1L inhibition is associated with reduced accessibility of intronic and intergenic enhancers that are bound *in-vivo* by the stemness-conferring transcription factor SOX2.

## RESULTS

### Multi-omics dataset reveals consistent epigenetic and transcriptional dynamics during ES-derived neuronal differentiation

To characterize the epigenetic and transcriptional changes during neuronal differentiation and to study the cell-type specific causal contribution of DOT1L to the neuronal differentiation process, we generate and integrate a multi-omics dataset encompassing comprehensive epigenomes of seven histone modifications (H3K4me1, H3K4me3, H3K9me3, H3K27ac, H3K27me3, H3K36me3, and H3K79me2) (in duplicates) and transcriptomes (triplicates) of mESC and NPC48h treated with dimethyl sulfoxide (DMSO) or Pinometostat (EPZ5676, EPZ). For NPC48h, we also generate chromatin accessibility profiles for each treatment regime in duplicates. To allow for a quantitative assessment of epigenetic changes, we use RELACS, a chromatin barcoding strategy for multiplexed and quantitative ChIP-seq (6) (Fig 1a).

**Figure 1.**
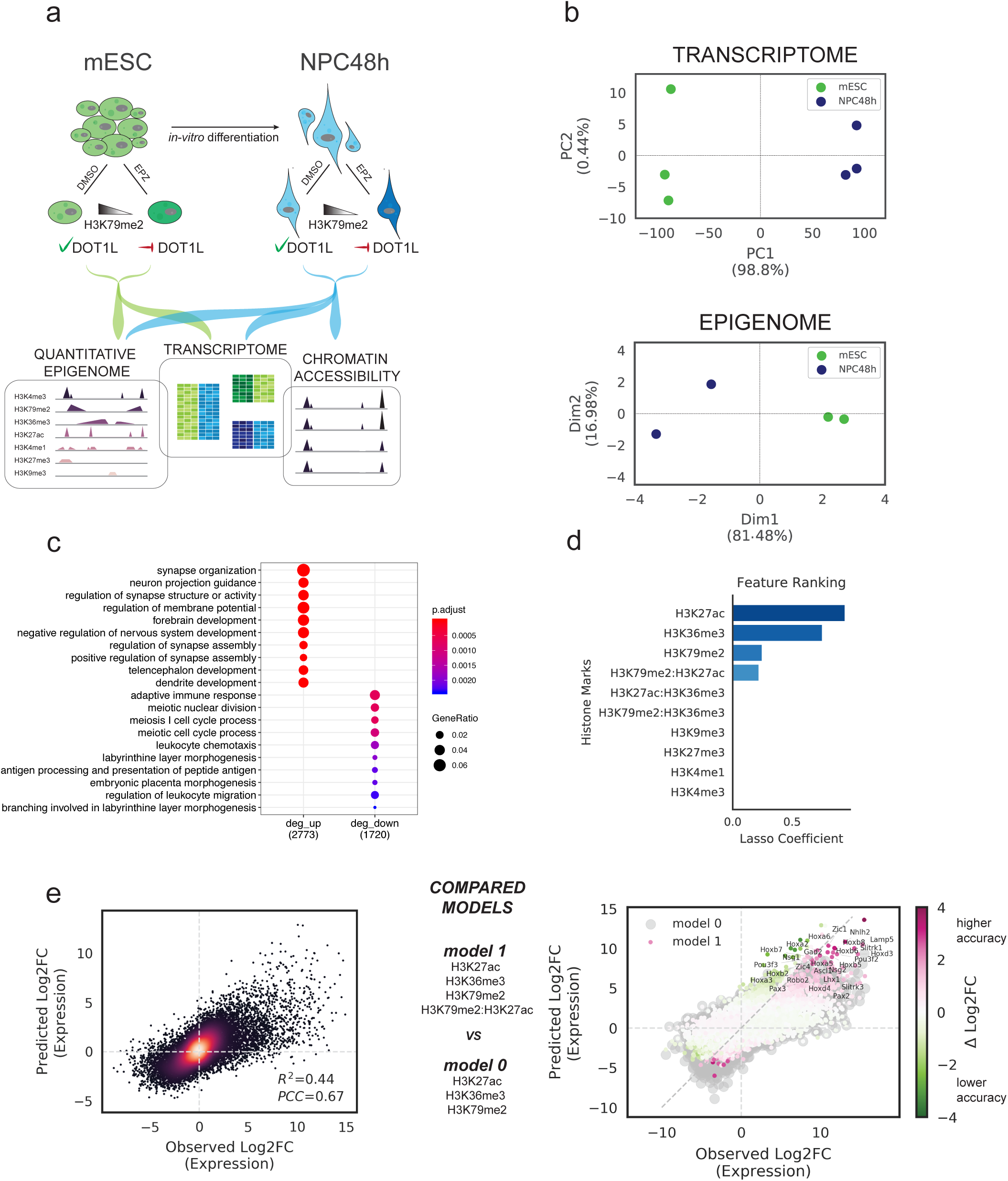
Multi-omics mapping and modelling of *in-vitro* neuronal differentiation. **a)** Experimental design of this study. We differentiate mESC towards neural progenitors (NPC48h) *in-vitro*. We treat mESC and NPC for 48h either with the DOT1L inhibitor EPZ5676 (10 nM) or with DMSO (1/1000) as control. For each sample, we generate transcriptomics data via RNA-seq and comprehensive epigenomes using quantitative ChIP-seq. We further map the accessible chromatin landscape for NPC48h using ATAC-seq. **b)** Upper panel: principal component analysis of the transcriptome of mESC and NPC48h on the top 500 most variable genes (rlog transformed counts) shows a separation between the two cell types on the first principal component. Lower panel: multiple factor analysis of the epigenome of mESC and NPC48h computed over the top 500 most variable 2kb windows for each histone modification yield similar results. Biological replicates are denoted by the same color. **c)** Top 10 most significant over-represented GO terms (adjusted p-value < 0.05) based on significantly upregulated (left) and downregulated (right) genes (abs(log2 fold-change) > 1, adjusted p-value < 0.01) in the comparison between ES-derived NPC48h and mESC. Genes increasing their expression in NPC48h are enriched for neuronal differentiation terms. **d)** Lasso regression coefficients are used to rank all input features. We retain a sparse model to predict transcriptional changes with three histone marks (H3K27ac, H3K36me3, H3K79me2) and an interaction term H3K79me2:H3K27ac. **e)** Fit of the multiple linear regression model. Observed vs predicted log2 fold-changes of gene expression (NPC48h vs mESC) as predicted through the linear combination of log2 fold-changes of H3K27ac, H3K36me3, H3K79me2 and H3K27ac:H3K79me2 (R^2^=0.44) (left panel). Predictions for different genes are differently affected by the interaction term (right panel). The biggest improvement in predictive accuracy is achieved for genes that are known targets of retinoic acid mediated neuronal differentiation (*Hoxa*, *Hoxb* cluster genes, *Ascl1, Zic1, Zic4, Pou3f2*, *Pou3f3*, *Nhlh2*, *Lhx1*).

We first assess the biological coherence of the generated multi-omics datasets. As expected, the transcriptome clearly segregates mESC and NPC48h into two distinct groups (Fig 1b, upper panel). A clear separation between mESC and NPC48h is also obtained from dimensionality reduction of the epigenome (Fig 1b, lower panel). The chromatin-based separation between cell types is most strongly determined by active histone modifications (Fig S1a). Differential gene expression analysis shows dynamic genes (abs(log2 fold-change) > 1, adjusted p-value < 0.01) to be prevalently upregulated in NPC48h compared to mESC (Fig S1b). Consistently, protein coding genes show higher coverage of the co-transcriptionally regulated marks H3K79me2 and H3K36me3 on the 5’end and 3’end of the gene body respectively, in NPC48h compared to mESC (Fig S1c). GO term enrichment analysis of differentially expressed genes (DEG) between NPC48h and mESC shows a clear neuronal signature in the upregulated set, providing evidence for the neuronal transcriptional identity of the differentiated cells (Fig 1c).

Next, we model the relationship between transcriptional dynamics and changes in histone modifications around transcriptional start sites (TSS) and transcriptional termination sites (TTS). As expected, chromatin changes correlate with transcriptional changes (Fig S1d), but the epigenetic features are collinear and thus redundant. To decrease this redundancy, we rank histone PTM dynamics based on their relevance for predicting transcriptional changes by fitting a regularized linear model to our dataset. We find that H3K27ac, H3K36me3 and H3K79me2 are selected as the most relevant predictive features for transcriptional dynamics, followed by an interaction term between H3K79me2 and H3K27ac (H3K79me2:H3K27ac) (Fig 1d). Our model (model 1, m_1) successfully captures the observed expression trends (R^2^ = 0.44) (Fig 1e, left panel) and results in a better fit compared to previous attempts with more complex models (28).

To confirm that the interaction H3K79me2:H3K27ac does not lead to overfitting, we compare model 1 to five alternative linear models and evaluate their performance based on the Bayesian information criterion (BIC) (Fig S1e). The inclusion of the interaction term does not lead to overfitting and increases the accuracy of log2 fold-change prediction for genes that are strongly upregulated during the differentiation process. Interestingly, many of the genes that are most affected by the interaction term are known targets of retinoic acid (RA) mediated *in-vitro* neuronal differentiation (e.g *Hoxa* and *Hoxb* clusters, *Ascl1, Zic1, Zic4, Pou3f2*, *Pou3f3*, *Nhlh2*, *Lhx1*) (29, 30) (Fig 1e, right panel).

Together, these data provide evidence for the coherence of the generated multi-omics dataset and show that *in-vitro* neuronal differentiation correlates with relative epigenetic and transcriptional activation. We show that gene expression changes can be predicted using a linear combination of a subset of histone modification changes (H3K27ac, H3K36me3, H3K79me2) and that the interaction between H3K27ac and H3K79me2 plays an important role to account for expression changes in RA target genes driving neuronal differentiation.

### H3K27ac and H3K79me2 undergo opposite global changes during *in-vitro* neuronal differentiation

The computation of histone modification changes in a classic ChIP-seq experiment imposes a per-sample normalization that prevents the detection of global shifts. In contrast, the RELACS barcoding strategy we employed allows for quantitative estimations of genome-wide global histone modification changes between samples. To assess the global dynamics of each histone PTM during the differentiation process, we estimate global scaling factors from sequencing data by computing pairwise ratios of input normalized read counts allocated to each sample after demultiplexing (6).

We observe that in NPC48h H3K4me3, H3K4me1 and H3K27me3 do not show detectable global deviations from the mESC reference level. H3K36me3 and H3K9me3 show a mild global increase. Strong global changes are instead observed for H3K27ac and H3K79me2 during neuronal differentiation, with the former decreasing by a factor of ∼ 2 (2.3 ± 0.12) and the latter increasing by a factor of ∼ 4 (3.9 ± 0.05) (Fig 2a).

**Figure 2.**
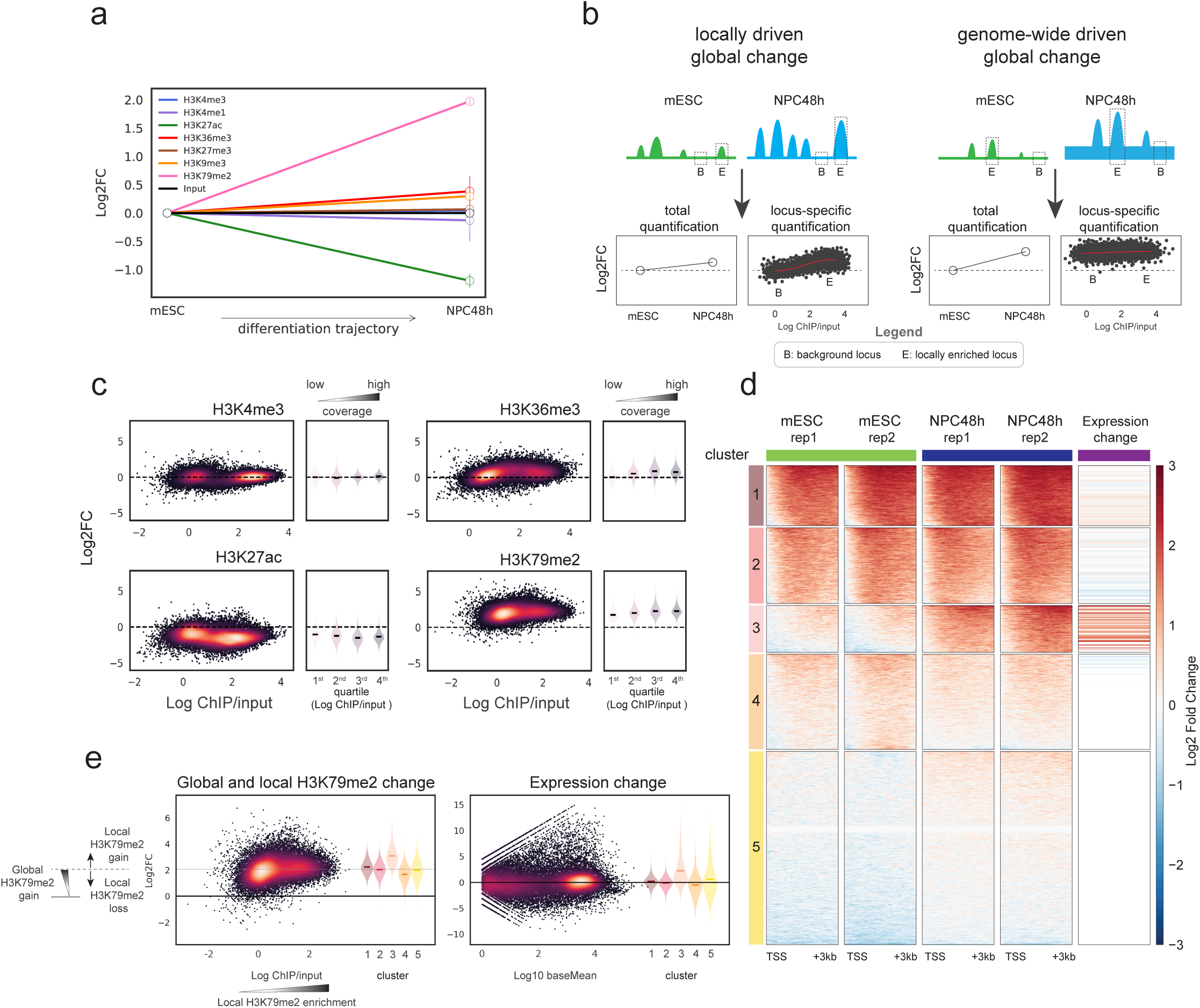
H3K27ac and H3K79me2 undergo opposite global changes that are independent of transcriptional changes, during *in-vitro* neuronal differentiation. **a)** Representation of the global scaling factors (log2 transformed) estimated in NPC48h using quantitative ChIP-seq with respect to the reference mESC level, for the seven histone modifications included in this study (n=2). Error bars denote one standard deviation from the mean. **b)** Model to illustrate that global histone modification changes can result from two different scenarios. Left panel: locally driven global changes may follow from increased number and/or magnitude of histone PTM local enrichment. Right panel: genome-wide driven global changes may follow from a genome-wide accumulation of a mark on both locally enriched regions (“peaks”) and on background regions. **c)** MA plots showing the mean coverage (x-axis) and log2 fold-change (y-axis) of four histone marks computed on bins overlapping annotated genomic features for the contrast NPC48h vs mESC (H3K4me3 and H3K27ac: transcription start site (TSS) ± 2kb; H3K79me2: TSS + 3kb; H3K36me3: transcription termination site (TTS) - 3kb). Next to each MA plot, we show a summary plot showing the global fold-change distribution (y-axis) for each quartile of mean coverage (x-axis). **d)** k-mean clustering (k=5) of H3K79me2 enrichment 3kb downstream of TSS of protein coding genes. A standard scaling method (by sequencing depth) and normalization (by input-control) was used. This highlights changes with respect to local background. The last column shows the log2 fold-changes in expression of the respective genes in each cluster for the contrast NPC48h vs mESC. Genes gaining local H3K79me2 enrichment tend to be upregulated during *in-vitro* neuronal differentiation. **e)** Left panel: MA plot showing the mean coverage (x-axis) and the global change in H3K79me2 (y-axis) computed on a window 3kb downstream of TSS of protein coding genes, next to a violin plot showing the global H3K79me2 changes for each of the 5 clusters previously identified in d). The scheme on the right helps interpretation of the global and local H3K79me2 changes. Right panel: MA plot of gene expression changes for the contrast NPC48h vs mESC, next to a violin plot showing the expression changes of genes clustered according to d).

Global shifts of histone PTM levels can be caused by two possible mechanisms. 1) Histone marks can accumulate on specific loci, resulting in local enrichment compared to flanking regions (so called “peaks”). A global shift can occur if the number and magnitude of histone PTM local enrichment changes across conditions. In this work, we refer to this mechanism as a *locally driven global change* (Fig 2b, left panel). 2) Alternatively, histone PTM can accumulate or be removed homogeneously over the whole genome, causing a base-level global gain/loss of the signal. Traditional ChIP-seq methods are unable to detect these global shifts. In this work, we refer to this mechanism as a *genome-wide driven global change* (Fig 2b, right panel). Eventually, global changes may result from a combination of 1) and 2).

To understand if the measured global changes are genome-wide or locally driven, we visualize locus-specific changes of H3K4me3, H3K27ac, H3K36me3 and H3K79me2 levels between NPC48h and mESC, on annotated genomic features (H3K4me3 and H3K27ac: transcription start site (TSS) ± 2kb; H3K79me2: TSS + 3kb; H3K36me3: transcription termination site (TTS) - 3kb). The results indicate that H3K4me3 levels are unaffected upon differentiation in both background and locally enriched regions. H3K36me3 levels do not change globally in background regions, but show a mild increase in locally enriched regions. This indicates that H3K36me3 global change is mostly locally driven. In contrast, loss of H3K27ac and gain of H3K79me2 affects background and locally enriched loci to an almost equal extent. This indicates that the global changes measured for these two marks are genome-wide driven (Fig 2c).

To provide a fully comprehensive picture of global histone modifications trends genome-wide, we perform chromatin segmentation, a method that reduces the high dimensionality of the epigenomic dataset by assigning a unique chromatin state attribute to each genomic bin based on the combination of histone modification enrichment (31, 32). For each histone mark, we compute log2 fold-changes between NPC48h and mESC over each of the 15 chromatin-state segments (E1 - E15). Extending our analyses to the whole genome and all chromatin states, the results confirm the trends computed over annotated gene bodies (Fig S2a), showing strong global changes only for H3K27ac and H3K79me2 during neuronal differentiation.

To explore candidate mechanisms accounting for global histone modification changes, we investigate the transcriptional dynamics of genes coding for epigenetic enzymes involved in the regulation of H3K27ac and H3K79me2. During differentiation, 4 of 13 expressed genes coding for proteins with histone deacetylase (HDAC) functions significantly increase their expression level (*Hdac9*, *Hdac11*, *Sirt2*, *Hdac2*) (log2 fold-change > 1, adjusted p-value < 0.01), while only 2 of 10 expressed genes coding for histone acetyltransferases (HAT) show significant changes in expression without consistent trend (*Kat6a*, *Hat1*) (Fig S1b). Assuming a proportional protein product, this observation suggests that the global decrease in acetylation may be partially driven by increased HDAC expression.

*Dot1l*, on the other hand, does not change its expression during the differentiation process. In yeast, Vos et al. (33) have shown that the grade (me1, me2, me3) and total level of methyl-H3K79 correlates with cell-cycle length and proliferation rate. To test whether global H3K79me2 differences estimated in our system are consistent with this model, we measure the proliferation rate of mESC and NPC48h. We find that mESC proliferate about 25 times faster compared to NPC48h (Fig S2b). The difference in proliferation rate between cell types qualitatively agree with the model advocated in Vos et al., but fails to account for the magnitude (4-fold increase) of the measured H3K79me2 difference.

In summary, we show that both H3K27ac and H3K79me2 levels change globally during *in-vitro* neuronal differentiation, with opposite trends. Both histone marks change through genome-wide acting mechanisms. The mechanism responsible for H3K79me2 global increase during differentiation still remains obscure, but we show that the proliferation rate alone does not suffice to account for the measured effect size.

### Local changes in H3K79me2 correlate with transcriptional activation

It has been reported that H3K79me2 local enrichment is the best linear predictor of gene expression levels (34), but the functional relevance of the global H3K79me2 increase during neuronal differentiation remains to be clarified, particularly in the context of transcription. Therefore, we ask whether differential H3K79me2 local enrichment, and/or the global H3K79me2 increase, associates with transcriptional dynamics.

To address this question, we stratify protein coding genes in 5 clusters using standard log2-ratio scores between sequencing-depth normalized H3K79me2 and input, individually for each cell type. Notice that this approach corresponds to a traditional normalization that absorbs all global changes. We quantify scores on a 3kb window downstream of TSS of mESC and NPC48h (Fig 2d). We observe that genes included in cluster 1 and 2 are locally enriched in both cell types and despite the global gain in H3K79me2, their expression levels do not change during differentiation (Fig 2e). Cluster 3, on the other hand, identifies a group of genes that gains H3K79me2 locally in addition to the global increase during development. These genes present a clear axonogenic signature (Fig S2c) and their expression level significantly increases during differentiation (Fig 2d, 2e). A mild reduction of H3K79me2 local enrichment is detected on genes belonging to cluster 4, but no major effect is observed at the transcriptional level. Eventually, cluster 5 identifies genes with neither H3K79me2 enrichment nor dynamic expression.

Together, these data show that, in our system, dynamic local enrichment of H3K79me2 associates with transcriptional activation (cluster 3) of genes critical for neuronal development. Global accumulation of H3K79me2 does not associate with transcriptional changes.

### Acute DOT1L inactivation is sufficient to bias the transcriptional state of NPC48h towards neuronal differentiation

H3K79me2 has been generally associated with transcriptional activity in yeast, fly, mouse and human. The mark is found in euchromatic regions and its enrichment strongly correlates with gene expression level (14). Yet, little is known about the causal relevance of DOT1L methyltransferase activity for the transcriptional process. To investigate the cell-type specific causal contribution of DOT1L enzymatic function for the genome-wide transcriptional activity and for the overall epigenetic context, we inhibit the enzyme in mESC and NPC by treating cells for 48 hours with the S-adenosyl methionine (SAM) competitor Pinometostat (EPZ5676, EPZ). Subsequently, we quantify transcriptional and epigenetic changes compared to cells treated with dimethyl sulfoxide (DMSO) as control.

To assess whether EPZ treatment successfully inhibited DOT1L methyltransferase activity, we compare H3K79me2 signal between treatment groups. Quantification after immunoblotting shows that the total H3K79me2 signal equals to 47.8% ± 5.7% and 59.6% ± 4.7% of the reference DMSO level in mESC and NPC48h respectively (Fig 3a). Quantification based on RELACS ChIP-seq confirms this trend and indicates that the total H3K79me2 signal is equal to 44.9% ± 2.4 % and 64.2% ± 5.8% of the reference DMSO level in mESC and NPC48h respectively (Fig 3a). In agreement with immunoblotting estimates, we observe that mESC lose more H3K79me2 compared to NPC48h. Cell-type specific differences are expected to occur as a consequence of unequal replication-dependent and independent histone turnover rate (Fig S2b) (35). To test if we could resolve signal loss at single-locus resolution, we compute locus specific changes of H3K79me2 over the previously defined five clusters (clustering analysis from Fig 2d). Results indicate that signal loss can be read as a function of H3K79me2 local enrichment (Fig 3b), where weakly marked loci (cluster 5) loose comparably less H3K79me2 signal than strongly marked loci (cluster 1).

**Figure 3.**
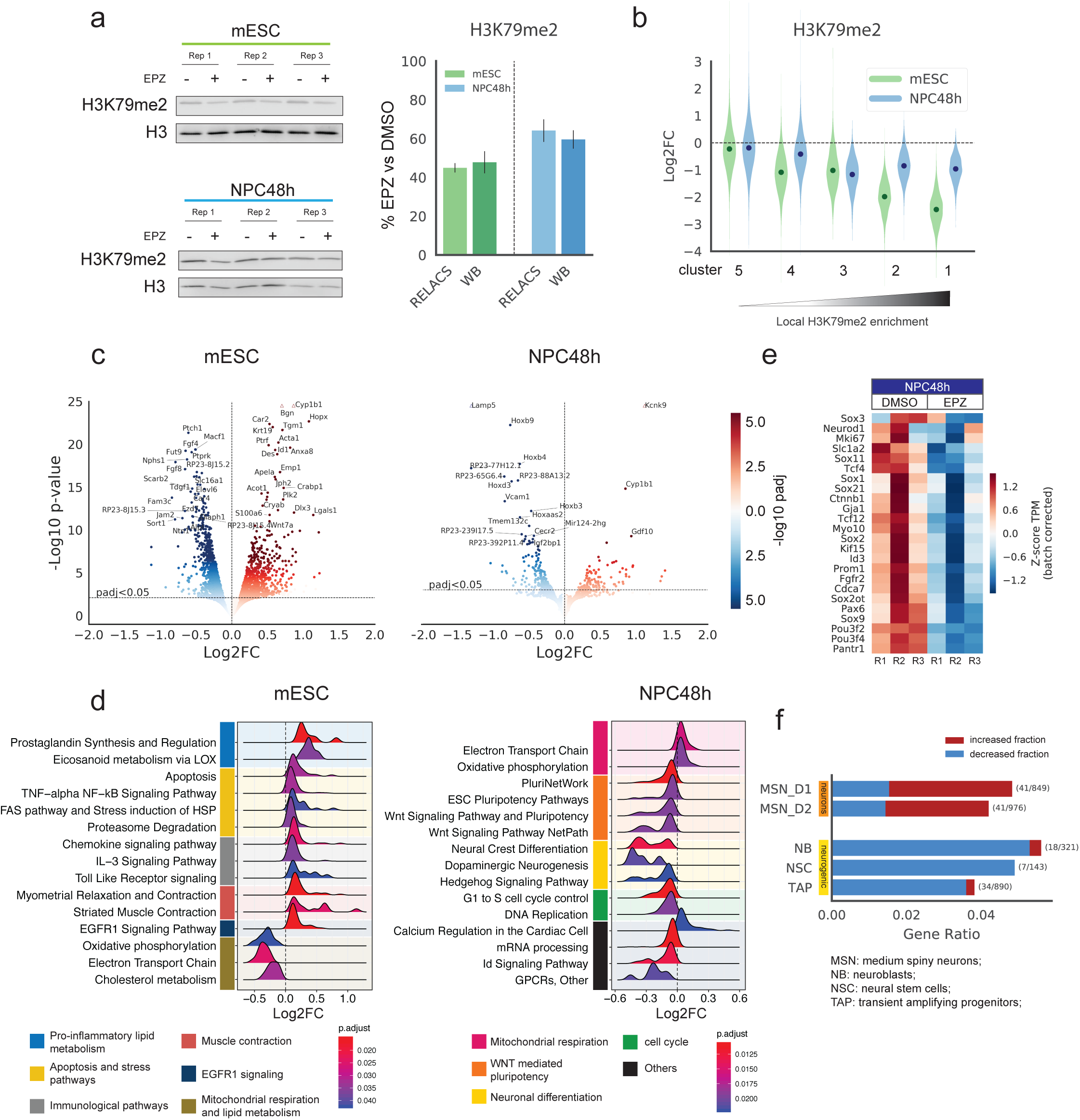
DOT1L inhibition for 48h alters the transcriptome of NPC towards neuronal differentiation. **a)** EPZ5676 treatment for 48h reduces the level of H3K79me2 in mESC and NPC48h. Left panel: immunoblotting of H3K79me2 and H3 (loading control) of EPZ-treated and DMSO-treated mESC (upper panel, green) and NPC48h (lower panel, blue). Right panel: barplot showing the global H3K79me2 signal for immunoblotting and RELACS ChIP-seq in EPZ treated mESC and NPC48h, represented as a fraction of the respective H3K79me2 level in DMSO treated cells. For RELACS data, we calculated the ratios of uniquely and high quality mapped reads (mapq > 5) and divided by the ratio for the respective inputs. **b)** Global fold-change of H3K79me2 in EPZ treated cells compared to DMSO, over the 5 clusters identified in the differentiation analysis of Figure 2d. High H3K79me2 local enrichment results in high loss upon EPZ treatment. **c)** EPZ treatment alters the transcriptome in mESC and NPC48h. Volcano plots showing log2 fold-change (x-axis) and -log10(p-value) (y-axis) of all genes tested for differential expression for the contrast EPZ vs DMSO treated mESC (left panel) and NPC48h (right panel). Genes are color-coded according to the respective adjusted p-value. Shade of blue is used for genes with decreased expression, while shade of red is used for genes with increased expression. The top most significant gene names are shown. **d)** Transcriptional deregulation induced by EPZ treatment affects groups of genes that are functionally coherent. Ridgeplot representing the expression log2 fold-change (EPZ vs DMSO) distribution (x-axis) of the leading edge genes from the top 15 most significant pathways (ranked on adjusted p-value), identified by running GSEA against the wikiPathways database, in mESC (left panel) and NPC48h (right panel). Each distribution is shaded according to adjusted p-value of the associated pathway. Pathways are grouped and color-coded according to functional similarity. **e)** EPZ treatment decreases expression of neural progenitor marker genes. Heatmap showing the z-score of batch corrected expression of various neuronal stem cell markers in DMSO and EPZ treated samples. Triplicates for each treatment group are shown. Expression values are normalized using transcripts per million (TPM). Positive z-scores are in shades of red, while negative ones are in shades of blue. **f)** Barplot showing the proportion of marker genes of three neurogenic cell types (NB, NSC, TAP) and two fully differentiated neuronal types (MSN_D1, MSN_D2) (36) that are differentially expressed in NPC48h following EPZ treatment (gene ratio, x-axis). Neurogenic marker genes are preferentially downregulated (blue), while marker genes of fully differentiated neurons are preferentially upregulated (red) upon EPZ treatment. Next to each bar, the gene ratio is explicitated as the number of marker genes that are differentially expressed in our dataset (numerator) over the total number of marker genes for each cell-type (denumerator).

To study the effects of DOT1L inhibition on the transcriptome, we first identify differentially expressed genes (DEG) across treatment groups. EPZ treatment causes a mild alteration of the transcriptome in both mESC and NPC48h, as indicated by principal component analysis and sample clustering on normalized count data (Fig S3a), where the main variability is from biological replicates rather than treatment. As a result, differentially expressed genes show only moderate log2 fold-changes (Fig 3c). Transcriptional alteration is more pronounced in mESC than NPC48h. 58 genes are differentially expressed in both cell types (adjusted p-value < 0.05). They follow a consistent fold-change trend, which may suggest a common underlying regulatory mechanism (Fig S3b).

Next, we identify annotated pathways and Gene Ontology (GO) terms associated with the transcriptional deregulation. For mESC, gene set enrichment analysis (GSEA) identifies significant pathways sharing an immunological and stress-induced pro-apoptotic molecular signature (Fig 3d, left panel). Among overrepresented gene ontology (GO) terms, we find increased expression of genes involved in actin cytoskeleton organization, and decreased expression of genes relevant for lipid and carbohydrate biosynthetic processes (Fig S3c, left panel). For NPC48h, GSEA shows deregulation of Wnt-mediated pluripotency pathways, neuronal differentiation and cell-cycle (Fig 3d, right panel), while among over-represented GO terms we find decreased expression of genes involved in embryonic organ development (e.g. *Hox* genes) and increased expression of genes coding for cation channels as well as genes involved in neuropeptides signaling pathway (Fig S3c, right panel).

DOT1L has been shown to prevent premature differentiation of the PAX6-positive neural progenitor pool in the developing cortex *in-vivo* (13). The functional signature observed in NPC48h suggests that acute DOT1L inhibition may be sufficient to induce a switch from a stemness-mediating to a differentiation-mediating transcriptional program. In line with this observation, we see a consistent decreased expression of a variety of neural stem cell markers in EPZ treated NPC48h (Fig 3e) (36, 37). To further substantiate this interpretation, we intersect our DEG set in NPC48h (EPZ vs DMSO treatment) with markers of neurogenic and neuronal cortical cell populations defined in two recent reports (36, 37). We find that differentially expressed markers of neurogenic cell populations, for the most part decrease in expression in our dataset, while differentially expressed markers expressed by fully differentiated neurons transcriptionally increase (Fig 3f).

Together, these results indicate that acute DOT1L inhibition for 48 hours is sufficient to deplete H3K79me2 on enriched loci genome-wide and to bring about mild yet functionally coherent transcriptional changes. Interpretation of the transcriptional response from a functional perspective suggests that DOT1L inhibition primes the transcriptome of NPC towards a neuronal differentiation stage.

### Acute DOT1L inhibition induces local epigenetic alterations linked to transciptional deregulation

The role of DOT1L as a chromatin writer demands a thorough analysis of the association between transcriptional and chromatin alterations. In mESC and NPC48h, quantitative ChIP-seq reveals that DOT1L inactivation does not consistently affect the global levels of histone modifications other than H3K79me2 (Fig 4a, S4a). Although EPZ treatment causes a decrease in H3K79me2 signal on every gene positively marked with this histone modification, the linear association of H3K79me2 depletion with transcriptional deregulation is weak in mESC (β = 0.027), and vanishingly small in NPC48h (β = 0.004) (Fig S4b). This indicates that acute DOT1L inhibition and subsequent reduction of H3K79me2 are not critical for immediate expression of most genes.

**Figure 4.**
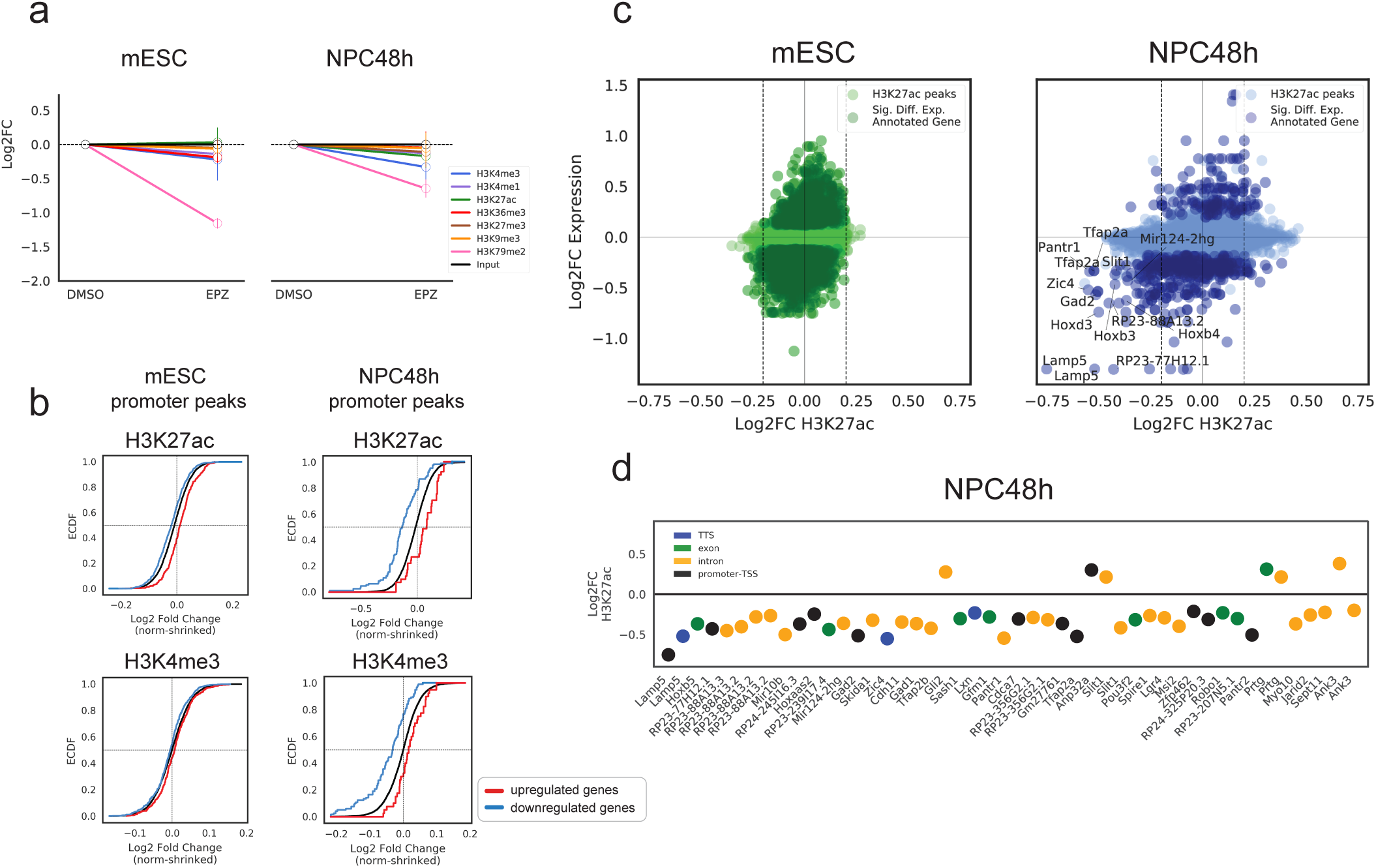
DOT1L inactivation results in local epigenetic changes that associate with transcriptional deregulation in NPC48h. **a)** Representation of log2 fold-changes of global scaling factors estimated via quantitative ChIP-seq in EPZ-treated cells with respect to the reference DMSO level, for the seven histone modifications included in this study. Left panel: global changes estimated in mESC. Right panel: global changes estimated in NPC48h. **b)** Promoter-associated active marks change consistently with EPZ-induced transcriptional dynamics. Upper panel: empirical cumulative density function (ECDF) of log2 fold-change of H3K27ac peaks overlapping promoter (TSS −1000bp, +500 bp) of differentially expressed genes (EPZ vs DMSO, adjusted p-value < 0.05) in mESC (top-left) and NPC48h (top-right). The red line shows the ECDF of log2 fold-change of H3K27ac peaks overlapping promoters of upregulated genes upon EPZ treatment, while the blue and black line depicts the same information for downregulated genes and all annotated genes respectively. Lower panel: Empirical cumulative density function (ECDF) of log2 fold-change of H3K4me3 peaks overlapping promoter of differentially expressed genes (EPZ vs DMSO, adjusted p-value < 0.05) in mESC (bottom-left) and NPC48h (bottom-right). The red line shows the ECDF of log2 fold-change of H3K4me3 peaks overlapping promoters of upregulated genes, while the blue and black line depicts the same information for downregulated genes and all annotated genes respectively. Genes that are transcriptionally affected as a consequence of EPZ treatment show a corresponding gain/loss of H3K27ac and H3K4me3 in their promoters. The epigenetic response is evident in NPC48h, while it is almost absent in mESC. **c)** H3K27ac peaks are depleted in a targeted set of genes in NPC48h upon DOT1L inhibition. Scatterplot showing the association between log2 fold-change of H3K27ac peaks (x-axis) and the expression log2 fold-change of annotated genes (y-axis) upon EPZ treatment for mESC (left, green) and NPC48h (right, blue). A peak is annotated to a gene if the peak overlaps any feature of the gene (promoter-TSS, introns, exons, TTS) or if it is proximal to the TSS/TTS (± 1kb). Each dot represents a H3K27ac peak. Darker dots represent H3K27ac peaks overlapping differentially expressed genes (adjusted p-value < 0.05) upon DOT1L inhibition. Peaks showing a significant loss of H3K27ac in NPC48h upon EPZ treatment are annotated with the gene symbol of the corresponding overlapping gene. **d)** Differential H3K27ac peaks annotated to differentially expressed genes upon DOT1L inhibition are preferentially found on intronic and promoter regions. Log2 fold-change of differential H3K27ac peaks (y-axis) overlapping or proximal to differentially expressed genes in NPC48h upon EPZ treatment. Each dot represents a H3K27ac peak. Peaks are colored based on the overlapping genomic feature (blue: TTS, green: exon, yellow: intron, black:promoter-TSS).

We observe, however, a difference in the mean expression level of genes that are transcriptionally affected upon EPZ treatment. Specifically, upregulated genes tend to be lowly expressed, while downregulated genes tend to be highly expressed (Fig S4c). H3K27ac correlates with expression level and recent studies suggested that H3K79me2 is important for maintaining H3K27ac enrichment on gene promoters and enhancers (25, 26). To verify whether H3K27ac signal is affected as a consequence of EPZ treatment, we perform differential analysis of H3K27ac peaks. Overall, we observe few significant changes in the profile of H3K27ac peaks upon EPZ treatment compared to the reference DMSO-treated samples, for both mESC and NPC48h. Log2 fold-change estimates of H3K27ac peaks overlapping the promoter of DEG show a weak trend consistent with expression changes, i.e. genes with increased expression tend to have higher levels of H3K27ac in promoter regions and vice versa (Fig 4b, upper panel). Notably, the effect size is stronger in NPC48h compared to mESC, despite a smaller number of genes being transcriptionally affected in the former cell type compared to the latter. A similar trend can also be observed for H3K4me3 (Fig 4b, lower panel).

Annotation of H3K27ac peaks to overlapping/proximal genes reveals a weak genome-wide correlation between acetylation and transcriptional changes (Pearson correlation coefficient = 0.19 and 0.16 in mESC and NPC48h respectively) (Fig 4c). We observe a more evident loss of H3K27ac signal in a subset of genes with decreased expression upon DOT1L inhibition in NPC48h (Fig 4c, right panel). Detailed genomic annotation of differential H3K27ac peaks overlapping transcriptionally downregulated genes in NPC48h upon EPZ treatment, shows a preferential distribution on intronic and promoter regions (Fig 4d).

Together, these data show that the genome-wide depletion of H3K79me2 does not result in a comparable global or local loss of H3K27ac, which argues against the hypothesis that H3K79me2 is generally critical to preserve H3K27ac from being targeted by deacetylase complexes in our model (25). Instead, our data show that local epigenetic changes of active marks (e.g. H3K27ac, H3K4me3) are directly linked to transcriptional changes, as indicated by the small effect size and the specific association with deregulated genes.

### Transcriptional alteration caused by DOT1L inhibition is associated with chromatin state signature of protein coding genes

To systematically investigate whether the altered transcriptional state is related to chromatin states, we use the chromatin segmentations of the control samples from mESC and NPC48h to measure the fraction of each chromatin state overlapping the promoter and the gene body of protein coding genes genome-wide. We apply t-distributed stochastic neighbour embedding (tSNE) to visualize the distribution of genes in a reduced 2-D space (38). Mapping of the DEG set reveals a clear separation between upregulated and downregulated genes, which is consistent across cell types (Fig 5a). Specifically, we observe that upon DOT1L inhibition, genes with a null, Polycomb repressed (H3K27me3) or bivalent (copresence of H3K4me3 and H3K27me3) promoter state are predominantly upregulated, while genes marked with an active promoter state (copresence of H3K4me3 and H3K27ac) tend to be downregulated (Fig 5a).

**Figure 5.**
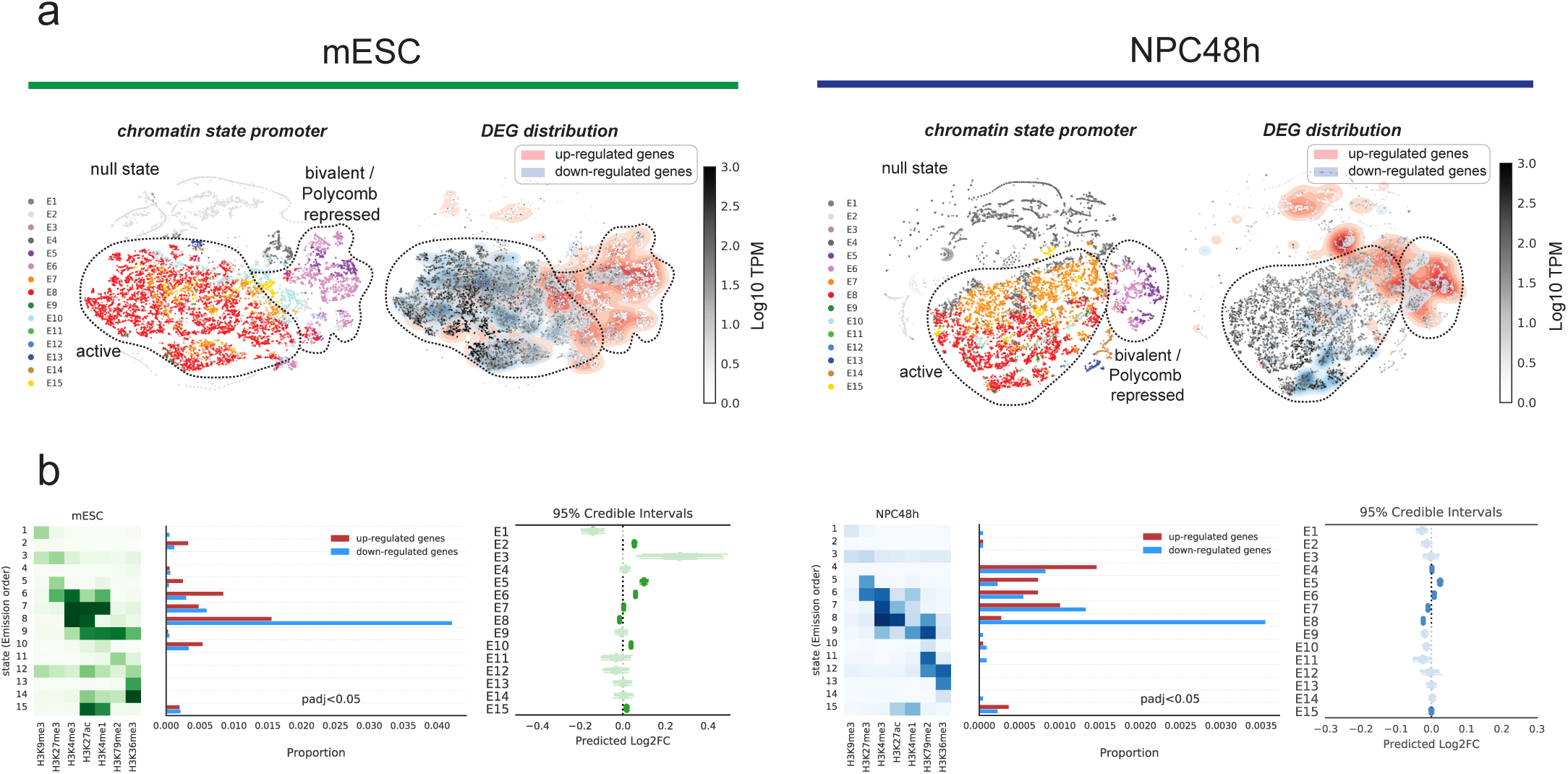
Transcriptional response to DOT1L inhibition associates with the chromatin state signature of protein coding genes. **a)** Dimensionality reduction (tSNE, perplexity = 30) of the chromatin state signature of protein coding genes for mESC (left, under the green stripe) and NPC48h (right, under the blue stripe). Genes are represented as dots; genes proximal to each other in the tSNE map have similar chromatin states fractions in their promoter and gene body. For each cell type, color-code is based on the most abundant chromatin state present in the promoter region on the left map, while on the right map, color-code is a gradient showing the expression level of each gene (Log TPM). Here, a 2D kernel density plot was over-imposed to show the distribution of differentially expressed genes (adjusted p-value < 0.05) on the tSNE map. **b)** Promoter chromatin state signature is weakly associated with transcriptional deregulation. For mESC (on the left, in green) and NPC48h (on the right, in blue) we show a heatmap summarizing the emission probability of the learned hidden markov model that was used to perform the chromatin segmentation, next to a histogram showing the proportion of differentially expressed genes (adjusted p-value < 0.05) classified according the the most abundant chromatin state present in their promoter region. Next to it, a plot showing the expected log2 fold-change posterior distribution (95% credible interval) of each group of genes sharing the same most represented chromatin state in the promoter, predicted via hierarchical bayesian modelling. Log2 fold-change posterior distributions of states that are not promoter-associated are shaded. For each group of genes sharing the same chromatin state in the promoter region, the expected mean log2 fold-changes in expression is estimated to be quite close to 0, suggesting that a small fraction of genes in each group is transcriptionally affected.

To quantify the strength of association between chromatin states and transcriptional deregulation, we fit a varying intercept model to estimate the expected transcriptional changes in each group of genes identified by the most represented chromatin state present in the promoter region. Results show that the estimated mean expression log2 fold-change in each gene group mildly deviates from 0, which suggests that the presence of any given chromatin state is not sufficient, per se, to induce transcriptional deregulation (Fig 5b).

Together, this evidence suggests that acute DOT1L inhibition results in derepression of silent genes and suppression of highly transcribed genes. Although the genes that are transcriptionally affected upon EPZ treatment are cell-type specific (Fig 3d, S3b), our analysis shows that they share a common epigenetic signature (Fig 5a), which hints towards a common underlying chromatin mechanism that could be shared across cell types. Despite there being an association between transcriptional deregulation and chromatin states, only a small subset of genes are affected by DOT1L inhibition. This prompts us to exclude that any specific combination of histone marks may be causally linked to the observed transcriptional phenotype. Instead, we hypothesize that the targeted transcriptional changes may be mediated by mistargeting of transcription factors (TF) regulating the subset of transcriptionally affected genes.

### DOT1L inhibition associates with decreased accessibility of a subset of intronic enhancers in NPC48h

To explore the hypothesis that DOT1L inactivation may affect specific DNA-binding TF, resulting in targeted gene expression changes, we focus on NPC48h, where the mild depletion of H3K79me2 allows to dissect the early response to H3K79me2 loss. To gain high resolution on putative transcription factor binding sites, we profile chromatin accessibility via ATAC-seq in NPC48h treated with DMSO or EPZ. We interrogate our data by taking a two-fold approach: on the one hand we identify associations between TF binding motifs and accessible promoter regions of DEG, and on the other we study how the accessibility profile is affected on enhancer regions genome-wide as a consequence of EPZ treatment (26).

To identify candidate TF associated with transcriptional alterations upon DOT1L inhibition in NPC48h, we identify motifs associated with accessible regions overlapping DEG promoters. Accessible promoter regions of genes that are upregulated upon EPZ treatment show high association with motifs bound by deacetylase complexes (i.e. SIN3A, HDAC2, REST) and basic Helix-Loop-Helix (bHLH) family members (i.e. ASCL1, NEUROD1, TCF21, TCF3) (Fig 6a, left panel). The enrichment of repressive complexes is consistent with the results of our chromatin-state analysis, which shows that promoters of upregulated genes are associated with null, Polycomb repressed (H3K27me3) or bivalent (H3K4me3 and H3K27me3) promoter states. Members of the bHLH family of TF are pivotal drivers of neuronal differentiation. In particular the proneural factor ASCL has been shown to direct neuronal cell fate specification by targeting repressed chromatin, acting as a pioneer factor, and to control the timing of neuronal differentiation (39, 40). Thus, these results support the hypothesis that acute DOT1L inhibition may be sufficient to initiate a specific transcriptional program towards neuronal differentiation.

**Figure 6.**
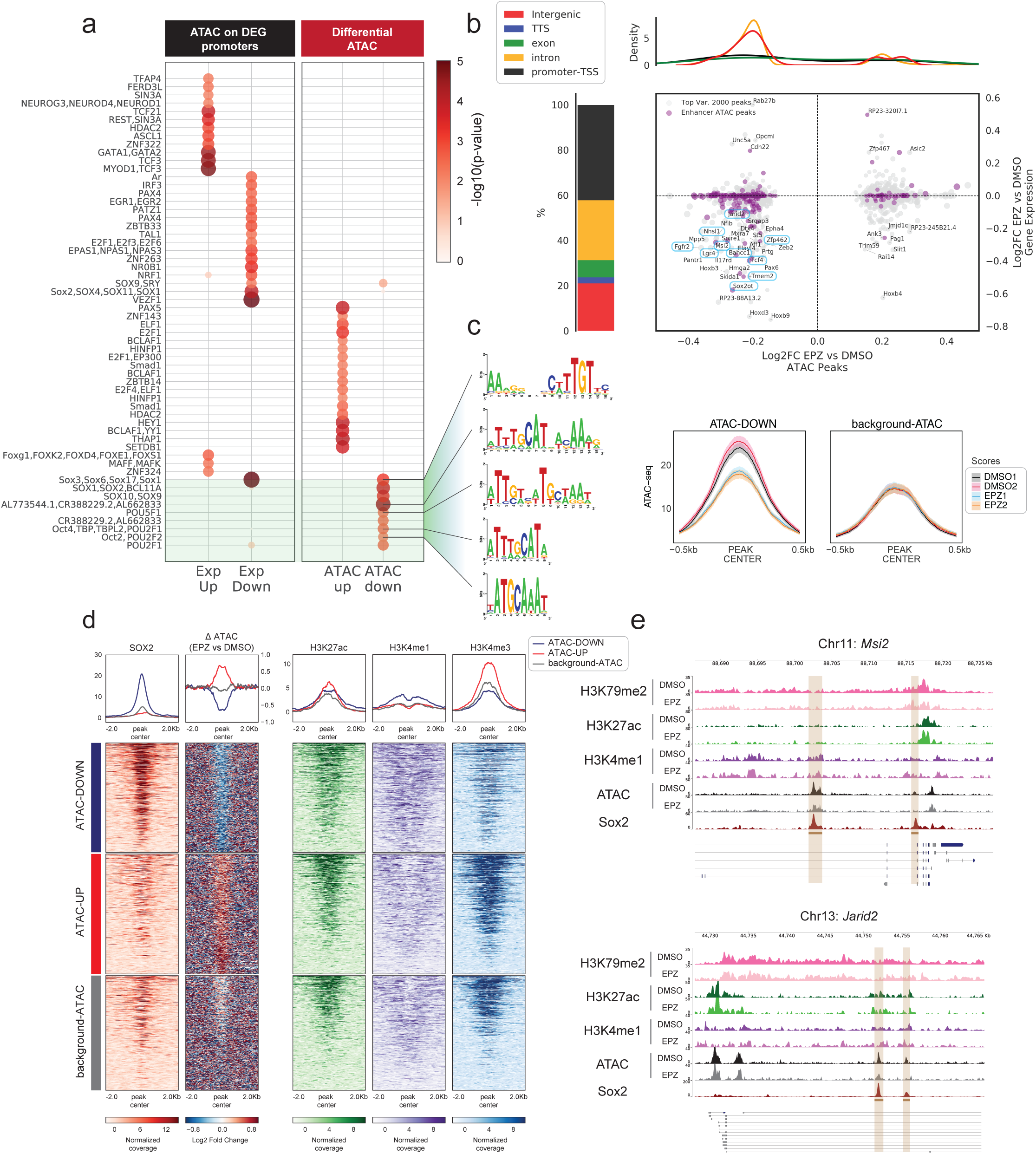
DOT1L inhibition decreases accessibility of SOX2 target loci in NPC48h. **a)** Left panel: differential motif analysis on ATAC peaks overlapping the promoter of differentially expressed genes upon EPZ treatment in NPC48h. Right panel: differential motif analysis on differentially accessible ATAC peaks upon EPZ treatment in NPC48h (right panel, under the red header). Size and color of each dot is proportional to -log10(p-value) associate with the motif. **b)** Characterization of dynamic ATAC peaks. Left panel: stacked barplot summarizing the genomic distribution of the 2000 most dynamic ATAC peaks (blue: TTS, green: exon, yellow: intron, black:promoter-TSS). Central panel: scatterplot showing the association between the log2 fold-change of dynamic ATAC peaks (x-axis) and the log2 fold-change of genes overlapping or proximal to at least one dynamic ATAC peak (y-axis) upon EPZ treatment in NPC48h. ATAC peaks overlapping enhancer regions are shown in purple. Gene symbols are shown for differentially expressed genes upon DOT1L inhibition. Genes that are associated with dynamic ATAC peaks and are downregulated in both mESC and NPC48h are highlighted in boxes. Top panel: density plot of the accessibility log2 fold-change of ATAC peaks overlapping enhancer upon EPZ treatment, stratified according to genomic annotation. Dynamic ATAC peaks overlapping enhancers tend to lose accessibility upon EPZ treatment and are found on intergenic and intronic regions. **c)** ATAC peaks with reduced accessibility upon DOT1L inhibition are associated with SOX/POU core motif. Left panel: logos of SOX and POU motifs showing highest association with ATAC peaks with reduced accessibility upon EPZ treatment in NPC48h. Right panel: metaprofile of ATAC-seq signal over ATAC peaks with reduced accessibility upon EPZ treatment (ATAC-Down, left subplot) and over random ATAC regions (background-ATAC, right subplot) for biological duplicates of DMSO and EPZ treated NPC48h. The prediction bands around the mean line show the 95% confidence interval. **d)** SOX2 preferentially binds *in-vivo* to regions with reduced accessibility upon EPZ treatment. Metaprofil and heatmap of SOX2 binding profile in brain-derived NPC (45) over open regions losing accessibility (ATAC-Down), gaining accessibility (ATAC-Up) and over unaffected regions (background-ATAC) upon EPZ treatment in NPC48h. Metaprofile and heatmap of the corresponding log2 ratio of EPZ vs DMSO ATAC-seq signal, H3K27ac, H3K4me1 and H3K4me3 input subtracted coverage of DMSO treated sample. **e)** Accessible loci with decreased accessibility upon DOT1L inhibition coincide with SOX2 binding in *Msi2* and *Jarid2.* Snapshots of *Msi2* (top panel) and *Jarid2* (bottom panel) loci, showing normalized coverage of H3K79me2, H3K27ac, H3K4me1, ATAC-seq for DMSO and EPZ treated NPC48h, and SOX2 coverage in brain-derived NPC (45). Highlighted regions show the genomic location of SOX2 peaks.

Open promoter regions of downregulated genes show enrichment for paired box (i.e PAX4) and SOX motifs, together with general GC rich motifs (Fig 6a, left panel). Our previous analyses have shown that downregulated genes upon DOT1L inhibition tend to be highly expressed and are associated with active promoter states (Fig S4c, 4d). Because highly expressed genes are often regulated by cell-type specific enhancers (41), we investigate the association between EPZ-induced transcriptional deregulation and enhancer activity.

Active enhancers are identified by the co-occurence of H3K27ac and H3K4me1 peaks and absence or low H3K4me3 coverage (2). Godfrey et al. have shown that accessibility of H3K79me2-marked enhancer and enhancer-associated H3K27ac decrease as a consequence of DOT1L inhibition (26). To study if accessibility of enhancer regions is perturbed in NPC48h upon DOT1L inhibition, we perform differential analysis of open chromatin regions between treatment groups. Similar to previous assays, we observe minor alterations of the accessibility landscape upon EPZ treatment, with very few regions reaching statistical significance as determined by DESeq2. Nevertheless, PCA identifies treatment regimes as the highest source of variance in the data (Fig S5a, left panel). To determine open chromatin regions with high contribution to the first principal component (PC1), we select 2000 peaks with the highest PC1 loadings, ranked on absolute value (Fig S5a, right panel), and we visualize the fold-change distribution of enhancer regions (Fig 6b, upper panel). The results show that intergenic and intronic enhancers tend to lose accessibility upon EPZ treatment. When we correlate dynamic accessible regions with expression changes of overlapping or proximal genes, we observe that loci with decreased accessibility are mostly associated with downregulated genes, regardless of enhancer status (Fig 6b, lower panel). Notably, this unbiased approach identifies 10 of only 39 genes that are commonly downregulated in both mESC and NPC48h upon DOT1L inhibition (*Jarid2*, *Fgfr2*, *Lgr4*, *Msi2*, *Bahcc1*, *Zfp462*, *Tcf4*, *Tmem2*, *Sox2ot, Nhsl1*), (3.15-fold enrichment, hypergeometric p-value = 0.00167) (Fig 6b, framed gene names, Fig S3b). Together, these observations suggest that decreased chromatin accessibility on enhancer regions in response to DOT1L inactivation, may contribute to the observed transcriptional downregulation.

### Intronic enhancer with decreased accessibility upon DOT1L inhibition are bound by SOX2 in brain-derived NPC *in-vivo*

To investigate whether decreased accessibility is specifically associated with the presence of H3K79me2, we measure H3K79me2 density on dynamic open chromatin regions (ATAC-Down, ATAC-Up) and on 1000 random open regions showing no change in accessibility as background (background-ATAC). Results show a clear association between H3K79me2 density and intronic open chromatin loci on ATAC-Down and background-ATAC regions compared to the other groups (two sided Mann-Whitney U test, p-value = 4.99·10^-7^) but no significant difference between intronic ATAC-Down and background-ATAC regions (two sided Mann-Whitney U test, p-value = 0.113) (Fig S5b). When we limit our study to open chromatin regions located over introns only, we see that 62% of protein coding genes having at least one ATAC peak with decreased accessibility are marked with H3K79me2, while only 25% and 34% of protein coding genes associated with ATAC-Up and background-ATAC regions are marked with H3K79me2 (Fig S5c).

Together, these data indicate that reduced chromatin accessibility upon DOT1L acute inhibition mostly, but not exclusively, affects regions marked with H3K79me2 in intronic loci. However, the presence of H3K79me2 alone - and its consequent loss upon EPZ treatment - is not a discriminant factor for decreased accessibility.

To study whether acetylation is altered as a consequence of EPZ treatment on dynamic ATAC regions, we visualize H3K27ac metaprofiles over ATAC-Down, ATAC-Up and background-ATAC regions, regardless of annotation class. On average, dynamic ATAC regions do not show any difference in H3K27ac levels (data not shown). To verify whether H3K27ac is affected on enhancers ATAC-Down depending on H3K79me2 presence (26), we visualize H3K27ac coverage on ATAC-Down enhancers high in H3K79me2 (H3K79me2 density > 45), on dynamic ATAC enhancers low in H3K79me2 (H3K79me2 density < 45) and on background-ATAC regions. Results indicate that H3K27ac is not selectively decreased on enhancers in ATAC-Down regions as a consequence of H3K79me2 presence (Fig S5d).

To evaluate whether regions with reduced accessibility are associated with a specific class of TF binding motif, we identify motifs associated with dynamic accessible regions. The analysis indicate that ATAC-Down regions present enrichment of POU/SOX core motifs (Fig 6c). This result is particularly interesting as SOX TF are critical regulators of neural progenitor pool maintenance and cell-fate specification (42–44). To verify that these loci are actually bound by SOX TF in neural progenitors *in-vivo*, we intersect the dynamic and background open chromatin regions identified in NPC48h upon EPZ treatment with publically available SOX2 ChIP-seq data generated on brain-derived neural progenitors (45). Metaprofile of SOX2 signal on dynamic ATAC-peaks and random background regions shows specific binding on open regions with decreased accessibility upon DOT1L inhibition (Fig 6d). Two exemplary loci, *Msi2* and *Jarid2*, show that decreased chromatin accessibility upon DOT1L inhibition coincide with regions bound by SOX2, local enrichment of H3K4me1/H3K27ac and presence of H3K79me2 (Fig 6e).

Together, these data indicate that, in our system, DOT1L inhibition results in decreased accessibility of a specific enhancer set s that is bound by SOX2 *in-vivo*. Loss of chromatin accessibility does neither associate with depletion of H3K27ac nor is it strictly correlated to co-occurence of H3K79me2.

## DISCUSSION

Here we report on a comprehensive multi-omics study of *in-vitro* neuronal differentiation and on the consequences of DOT1L inhibition for the differentiation process. This includes the study of the quantitative dynamics of chromatin modifications during *in-vitro* neuronal differentiation by use of a quantitative and high-throughput ChIP-seq method. This is, to our knowledge, the first application of a quantitative strategy to a physiological differentiation setting, and it reveals that the epigenome of neuronal committed cells undergoes global histone modification changes with respect to the pluripotent precursor.

Various studies have documented a progressive chromatin condensation during mESC differentiation (5,46,47), but contrasting evidence has been collected regarding the extent and relevance of global histone modification changes for cellular differentiation. For example, Ugarte et al. describe a progressive decrease in nuclease sensitivity during hematopoietic differentiation but fail to detect any significant global changes in histone modifications levels through immunoblotting assessing H3K4me3, H3K27ac, H3K16ac, H4K20me1, H3K36me3, H3K27me3, H3K9me2 and H3K9me3 (46). Efroni et al. characterize global transcriptional and epigenetic changes during mESC-derived NPC differentiation. Their evidence, based on immunoblotting and imaging, suggests that both global RNA levels and active histone modification abundances (e.g H3K4me3) are decreased in differentiated cells compared to the embryonic precursor (5).

In contrast to these previous studies, here we use a quantitative ChIP-seq protocol (RELACS) to estimate global histone modification changes during *in-vitro* neuronal differentiation. We find that only H3K27ac and H3K79me2 levels change globally, in opposite directions, during *in-vitro* neuronal differentiation, through genome-wide acting mechanisms.

Biologically, these results are notable in various respects. First, they suggest that the progressive chromatin condensation observed during *in-vitro* neuronal differentiation mostly follows from a genome-wide deacetylation process, while a smaller contribution may come from the global accumulation of repressive histone modifications (e.g H3K9me3) (48). Loss of H3K27ac is consistent with chromatin condensation (2), as H3K27ac is a mark associated with loose chromatin packaging and is known to be highly abundant in mESC (48, 49).

Secondly, our data show that H3K79me2 increases globally during neuronal differentiation *in-vitro*. We show that developmental gain of local enrichment of H3K79me2 associates with transcriptional activation of genes critical for neuronal differentiation. In contrast, global accumulation of H3K79me2 does not generally correlate with transcriptional activity. Our data suggest that global differences in H3K79me2, as measured during differentiation and as a consequence to pharmacological inhibition of DOT1L, may be partly caused by the different proliferation rates of mESC and NPC48h, in accordance with previous reports (33).

As a consequence of H3K79me2 global increase during *in-vitro* neuronal differentiation, we investigate the relevance of DOT1L methyltransferase activity for the establishment of chromatin and transcriptional states genome-wide in mESC and their differentiated progeny, NPC48h. The third main result of this study shows that DOT1L inactivation affects gene expression in a targeted manner, despite the genome-wide depletion of H3K79me2. Our results clearly indicate that the presence of H3K79me2 is neither generally critical for the deposition of other histone modifications, nor is it necessary for sustaining the expression levels of most genes. We observe depletion of H3K27ac upon EPZ treatment, which does not follow the global decrease in H3K79me2. Locally, however, loss of H3K27ac on enhancers and promoters alike correlates with transcriptional downregulation, and it is mirrored by a corresponding decrease in H3K4me3 on promoters.

Most importantly, we show that upon DOT1L inactivation, transcriptionally deregulated genes present a coherent chromatin signature in their promoter. Our data indicate that DOT1L inactivation associates with upregulation of genes with a repressed, poised or null promoter state, and downregulation of highly expressed genes marked with active histone modifications. Based on the data presented in this work, it is tempting to hypothesize that the transcriptional upregulation upon DOT1L inhibition observed in the mammalian system may result from impaired targeting of the chromatin by repressive complexes. The cause of this may either reside in the altered H3K79me distribution, as in the yeast model, or it may indirectly follow from the selective downregulation of highly transcribed genes coding for repressive proteins (e.g *Jarid2, Zfp462*) (50–53).

Finally, our study supports the view that the targeted transcriptional response to DOT1L inactivation may in part be explained by decreased accessibility of active enhancers bound by critical TF. Whereas in NPC48h, DOT1L inhibition results in decreased accessibility at chromatin regions bound by SOX2 *in-vivo*, the reduced chromatin accessibility is not accompanied by depletion of H3K27ac. In this light, our data partly contrasts with the model advocated by Godfrey at al. (26), which establishes a causal link between presence of H3K79me domains, preservation of H3K79me-rich enhancer activity and H3K27ac levels. Godfrey et al. have recently identified a class of enhancers dependent on H3K79 methylation, where the frequency of enhancer-promoter interaction is disrupted upon DOT1L pharmacological inhibition (26). Consistent with this report, we observe that EPZ treatment induces a loss in accessibility of a subset of intronic and intergenic enhancers. Although we find that decreased accessibility is associated with H3K79me2 enrichment in intronic open loci, our data also suggest that H3K79me2 enrichment is generically present over intronic ATAC peaks and does not discriminate between dynamic and non-dynamic open regions. Moreover, around 40% of intronic enhancers with decreased accessibility upon DOT1L inhibition are not strongly marked by H3K79me2. Together, our data indicate that DOT1L inhibition may alter the cellular transcriptional state by affecting only a subclass of H3K79me2-positive enhancers.

In conclusion, our findings agree with the model proposed by Godfrey et al. in that DOT1L inhibition results in decreased accessibility of H3K79me2-positive enhancers. In our system, though, we observe a specific response that pertains only to a subset of regulatory regions bound by sequence-specific transcription factors (e.g. SOX/POU). The closure of these cis-acting enhancers may be responsible for the transcriptional decrease of highly expressed, cell-type specific genes conferring stemness to progenitors. In addition, we here present first evidence explaining transcriptional increase upon DOT1L inhibition. We hypothesize that decreased expression of cell-type specific transcripts coding for proteins with repressive functions (e.g *Jarid2, Zfp462*), together with altered accessibility for deacetylation complexes, may result in derepression of silent genes localized on facultative heterochromatic regions.

## MATERIALS and METHODS

### mESC culture and *in-vitro* neuronal differentiation

mESC were cultured on inactivated MEF for 3 passages (p3) and from p4 onward on gelatin-coated plates (medium: 82% DMEM (Thermo Fisher, US), 15% FBS (Thermo Fisher), 1% Glutamax (Thermo Fisher), 1% PSN (Thermo Fisher), 1% NEAA (Thermo Fisher) + LIF (Sigma) (dilution = 1/1000) + β-Mercapto-EtOH (Thermo Fisher) (dilution = 1/500)). Feeder-free mESC were treated with either EPZ5676 (Hycultec) (10 nM), or DMSO (Thermo Fisher) (dilution = 1/1000) for 48h.

mESC were differentiated *in-vitro* towards NPC48h according to Bibel et al. (27). Briefly, feeder-free mESC were trypsinized and dissociated to create a single cell suspension. Cells were used to form cell aggregates (CA) on non-adherent (Grunier) plate (4·10^6^ single cells per plate; medium: 87% DMEM, 10% FBS, 1% Glutamax, 1% PSN, 1% NEAA + β-Mercapto-EtOH (1/500)). 4 days after CA formation, CA were exposed to retinoic acid (7.5 µM) for 4 days. CA were dissociated into single cells and seeded on PORN/LAMININ coated 6 well plates and grown in N2 medium for neuronal differentiation. At this stage, cells were treated either with EPZ5676 (10 nM) or DMSO (1/1000) for 48h. At treatment completion, NPC48h were collected for downstream processing.

### RELACS ChIP-seq

RELACS protocol was carried out according to Arrigoni et al. (6). Cells were fixed in 1% formaldehyde for 15 minutes. Reaction was quenched with 125 mM glycine for 5 minutes, followed by 2 washings with DPBS + proteinase inhibitor cocktail. Cell nuclei were isolated following Nexson protocol (54) and permeabilized with 0.5% SDS. Chromatin was digested *in situ* using restriction enzyme CviKI-1 (NEB, R0710L) and barcoded using RELACS custom barcodes (4bp UMI + 8bp RELACS barcode, see Table 1 for details). Nuclei from each sample were burst via sonication to extract the barcoded chromatin fragments and pooled into a unique tube. A single immunoprecipitation (IP) reaction for all samples included in this study was carried out on IP-star according to (6) (see Table 2 for antibodies details). Immunoprecipitated chromatin was used for Illumina library preparation (NEBNext Ultra II DNA Library Prep Kit) and sequenced on HiSeq 3000 Illumina machine (paired-end, read length 75 bp).

**Table 1:** RELACS custom barcodes

| Sample | RELACS barcode |
| --- | --- |
| mESC_DMSO_rep1 | TTCGCTCT |
| mESC_DMSO_rep2 | ACGTGTAC |
| mESC_EPZ_rep1 | TACCGATG |
| mESC_EPZ_rep2 | TTGGTTGG |
| NPC48h_DMSO_rep1 | CCTCTCAA |
| NPC48h_DMSO_rep2 | TTGTGGCT |
| NPC48h_EPZ_rep1 | CCGAATAC |
| NPC48h_EPZ_rep2 | TGTGATCG |

**Table 2:** Antibody details

| Histone modification | Ab details (company, product ID, lot) |
| --- | --- |
| H3K27ac | Diagenode, C1541096, lot A1723-041D |
| H3K27me3 | Diagenode, C15410195, lot A1811-001P |
| H3K36me3 | Diagenode, C15410192, lot A1847-001P |
| H3K4me1 | Diagenode, C15410194, lot A1863-001D |
| H3K4me3 | Diagenode, C15410003, lot A5051-001P |
| H3K79me2 | Abcam, ab3594 |
| H3K9me3 | Diagenode, C15410193, lot A1671-001P |

### RNA-seq

RNA was extracted using RNAeasy Mini Kit (Qiagen). Libraries were generated using the NEBNext Ultra RNA Library Prep Kit, following manual’s instructions. Libraries were sequenced on a HiSeq 3000 Illumina machine (paired-end, read length 150 bp).

### ATAC-seq

ATAC-seq libraries were generated according to (55). Briefly, ∼ 50.000 cells were washed in ice-cold PBS and incubated in transposition reaction mix (Nextera DNA Sample Preparation Kit). Transposed DNA was purified (MiniElute Kit, Qiagen) and PCR amplified for 5 cycles. We determined the number of additional PCR cycles via qPCR according to (55). Libraries were sequenced on a HiSeq 3000 Illumina machine (paired-end, read length 75 bp)

### Proliferation assay

Proliferation assay was performed using Click-iT® EdU Alexa Fluor 488 Flow Cytometry Assay Kits (C10425), following the manufacturer’s instructions. Intact nuclei were further stained with DAPI and analyzed on a BD LSRFortessa cell analyzer using BD FACSDiva software.

### Immunoblotting

mESC or NPC48h were lysed in RIPA buffer (1% NP-40, 1% SDS, 0.5% sodium deoxycholate diluted in Phosphate Buffered Saline, PBS). Cells were centrifuged (10 min, 13000 rpm) and the supernatant collected. Protein concentrations were determined with Bradford reagent (Bio-Rad). 15 µg of protein extract were loaded with 5x Laemmli buffer on Mini Protean TGX gels (Bio-Rad) and run at 100V for 1.5 h. Proteins were transferred to PVDF membranes (Trans-blot Turbo Transfer Pack) using the Trans-blot Turbo Transfer System (both from Bio-Rad) following manufacturer’s instructions. Membranes were blocked with 5% BSA in TBS-T (blocking buffer) for 1 h and incubated overnight with primary antibodies (diluted in blocking buffer). Membranes were washed, incubated with secondary antibodies for 1 h and detected using ECL or Femto substrates (Thermo Scientific) and LAS ImageQuant System (GE Healthcare, Little Chalfont, UK). The following antibodies were used: anti-H3K79me2 (1:1000 dilution, see Table 1 for details). For densitometric analyses, ImageJ software was used (56).

### Bioinformatics analysis

All sequencing data were processed with snakePipes (v. 1.1.1) (57). Relevant parameters used for each experiment and summary QC are available at https://github.com/FrancescoFerrari88/code_DOT1L_paper/tree/master/multiQC_ConfigParameters. Mapping was performed on mouse genome build mm10 (GRCm38). For ChIP-Seq and ATAC-seq, high quality and uniquely mapping reads were retained (mapq > 5). RELACS custom barcodes were designed with integrated UMI, so duplicate removal was performed using UMITools (58), while a standard deduplication was applied for ATAC-seq reads. We use gencode M18 as reference gene model throughout all analysis. For ChIP-seq and ATAC-seq data, snakePipes also provided candidate peak regions using MACS2 (default parameters).

Differential analysis for RNA-seq was carried out using DESeq2 (v. 1.22.1) (59) on count matrices output from snakePipes (featureCounts, v. 1.6.4). We used a linear model controlling for batch effects (e.g. ∼ *batch* + *treatment*) and we applied apeglm log2 fold-change shrinkage (60). We estimate fold-changes for each histone modification on annotated genomic features known to associate with local histone PTM enrichment (H3K4me3, H3K27ac, H3K4me1: narrow promoter (TSS ±1kb); H3K79me2, H3K27me3, H3K9me3: extended promoter (TSS −1kb,+3kb); H3K36me3: transcription termination site (TTS - 3kb,+0.5kb)).

Global differential ChIP-seq analysis was carried out after applying RELACS specific normalization by computing empirical log-fold changes across conditions (see “RELACS normalization and estimation of global histone modification changes”). Traditional differential ChIP-seq and ATAC analysis was performed on consensus peak sets, coverage was computed using deepTools’ multiBamSummary (61) and differential regions identified via DESeq2. We eventually applied normal log2 fold-change shrinkage. Peaks were annotated using Homer (v. 4.10) (62) and UROPA (v. 3.1.0) (63). We use GimmeMotifs (v. 0.13.1) for motif enrichment and differential motif analysis (64). Metaprofile of ChIP-seq and ATAC-seq signals were generated with deeptools (61) and deepStats (65).

Multiple factor analysis was done using FactoMineR (v. 1.41) (66). The algorithm was run on a matrix of shape 4 (samples) x 3500 (features). As features, we included the top 500 most variable 2kb loci for each of the seven histone modifications (feature groups), selected after applying variance stabilizing transformation to the counts matrix. We used scikit-learn (Python module) (v. 0.19.1) for principal component analysis and tSNE, while linear modeling was performed using sklearn and statsmodels (v. 0.9.0). GO enrichment analysis and pathway analysis were performed using clusterProfiler (v. 3.10.1) (67).

We used ChromHMM (31) with default parameters for chromatin segmentation. We trained two independent models for each cell type on the DMSO treated samples. We then used these models to perform the segmentation in the respective cell types for both treatments (EPZ and DMSO).

We compute the chromatin state signature of protein coding genes in mESC and NPC48h according to (38). For each gene, we identify potentially used transcripts by intersecting annotated TSS with H3K4me3 peaks. If a gene does not overlap with any H3K4me3 peak, we consider the full gene annotation. For each candidate gene, we then compute the fraction of overlap between each chromatin state segment in the control sample with the promoter region (TSS −1kb, +500 bp) and with the full gene body. In this way, each gene is identified by a vector of length 30 (15 states for the promoter + 15 states for the gene body). A matrix of shape g (number of genes per cell) x 30 is eventually used for dimensionality reduction by applying tSNE (68).

To compute the enrichment for the frequency of transition of each chromatin state in DMSO to each chromatin state in EPZ (Fig S4a), we first flatten the chromatin state segmentation across all samples. Next, we compute the frequency of transition across chromatin states from DMSO to EPZ. A transition is identified if it is concordant across replicates (foreground transition matrix). The background frequency (transition noise) is computed as the frequency of transition across chromatin states from DMSO_rep1 to DMSO_rep2 and from EPZ_rep1 to EPZ_rep2 (background transition matrix). The ratio between the foreground and background transition matrix results in the enrichment score.

Bayesian linear modeling was performed using pymc3 (v. 3.6) (69). The expected log2 fold-change for each group of genes (i) identified by the most represented chromatin state present in the promoter regions was identified by fitting the following hierarchical linear model:

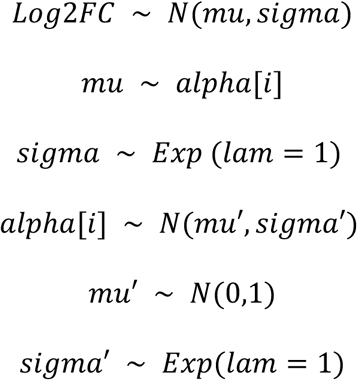

All visualizations were generated in Python (v. 3.6) and R (v. 3.5).

### RELACS normalization and estimation of global histone modification changes

To estimate global histone modification changes, first we demultiplexed fastq files on RELACS custom barcodes. Then, for each sample, we divided the number of uniquely and high-quality mapped read-pairs (mapq > 5) coming from a ChIP of interest by the total number of read-pairs coming from the respective input. For estimating global histone modification changes, we considered either the total number of mapped reads genome-wide. Pairwise quantitative comparisons between samples were computed as log2 ratio between input-normalized total mapped read counts.

Local changes were estimated in the same way, by repeating this procedure for each individual bin of interest.

### Data and code availability

The fully reproducible and documented analysis is available on github at <u>github.com/FrancescoFerrari88/code_DOT1L_paper</u>, as Jupyter notebooks and R scripts. Raw data and normalized bigWig tracks were deposited to GEO and are available for download using the following accession number: GSE135318.

## ACKNOWLEDGEMENTS

We would like to thank Ulrike Bönisch, Chiara Bella, Katrin Groβer and Steffen Wolter for providing essential support for the generation of the sequencing data. We thank Yaarub Musa and Gerhard Mttler for the fruitful discussions on the research project. Special thanks to Devon Ryan and Leily Rabbani for providing the scripts needed for RELACS demultiplexing and for integrating the RELACS workflow into the publically available NGS processing pipeline (snakePipes). We thank Alejandro Villarreal for help with mESC and NPC48h cultures and ATAC-seq. This research was funded by the Deutsche Forschungsgemeinschaft (DFG, German Research Foundation): 322977937/GRK2344 and SFB 992 (Medical Epigenetics).

---

**Supplementary 1.**
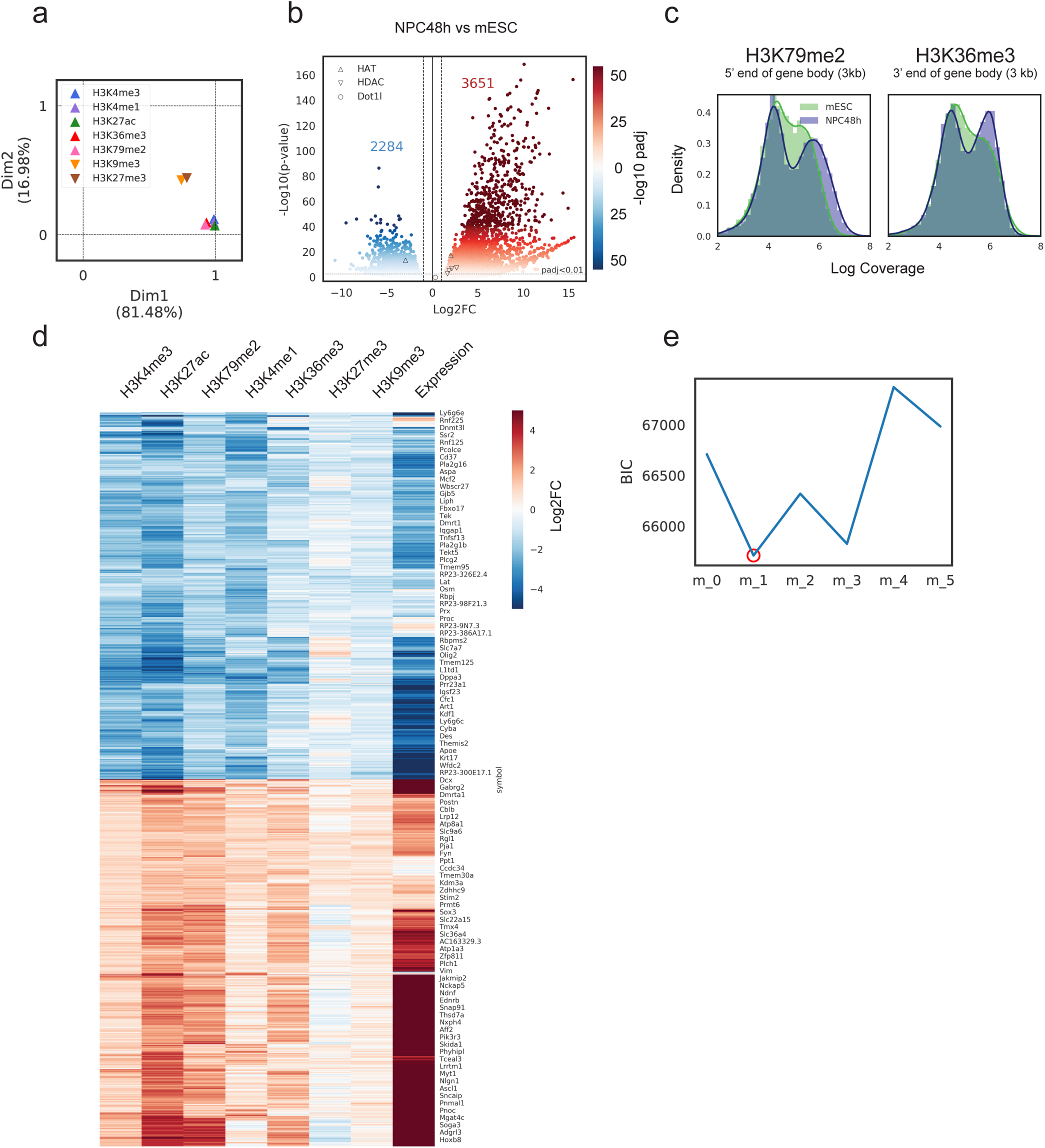
Neuronal differentiation correlates with relative transcriptional and epigenetic activation. **a)** Graph showing the contribution of each group of features (most variable 2kb bins for each histone modification) to the first and second dimension of the multiple factor analysis in Figure 1b, lower panel. **b)** Volcano plot summarizing differential expression analysis for the contrast NPC48h vs mESC. Genes are color-coded according to the -log10(adjusted p-value), genes with decreased expression in blue and with increased expression in red. HDAC: histone deacetylases; HAT: histone acetyltransferase. The dotted lines denote the thresholds used in this study (abs(log2 fold-change)>1, adjusted p-value<0.01). **c)** Distribution of H3K79me2 and H3K36me3 normalized coverage (relative log expression (RLE) normalization on background regions), computed over 3kb window downstream of TSS of protein coding genes and 3kb upstream of TTS of protein coding genes respectively, in mESC (green) and NPC48h (blue). TSS: transcription start site, TTS: transcription termination site. **d)** Heatmap showing the log2 fold-change (NPC48h vs mESC) of histone modifications (H3K4me3, H3K27ac, H3K4me1: narrow promoter (TSS ±1kb); H3K79me2, H3K27me3, H3K9me3: extended promoter (TSS −1kb,+3kb); H3K36me3: transcription termination site (TTS - 3kb,+0.5kb)) and the log2 fold-change in expression (NPC48h vs mESC) of the corresponding gene. We show the top 1000 genes with increased and decreased expression. **e)** Model selection. We employ the Bayesian information criteria (BIC) to select the model with minimal BIC score among 5 multiple linear regression models with different complexity (m_0: H3K27ac, H3K36me3, H3K79me2; m_1: H3K27ac, H3K36me3, H3K79me2, H3K79me2:H3K27ac, m_2: H3K27ac, H3K36me3, H3K79me2, H3K9me3, H3K27me3, H3K4me1, H3K4me3, m_3: H3K27ac, H3K36me3, H3K79me2:H3K27ac; m_4: H3K36me3, H3K79me2; m_5: H3K27ac, H3K36me3).

**Supplementary 2.**
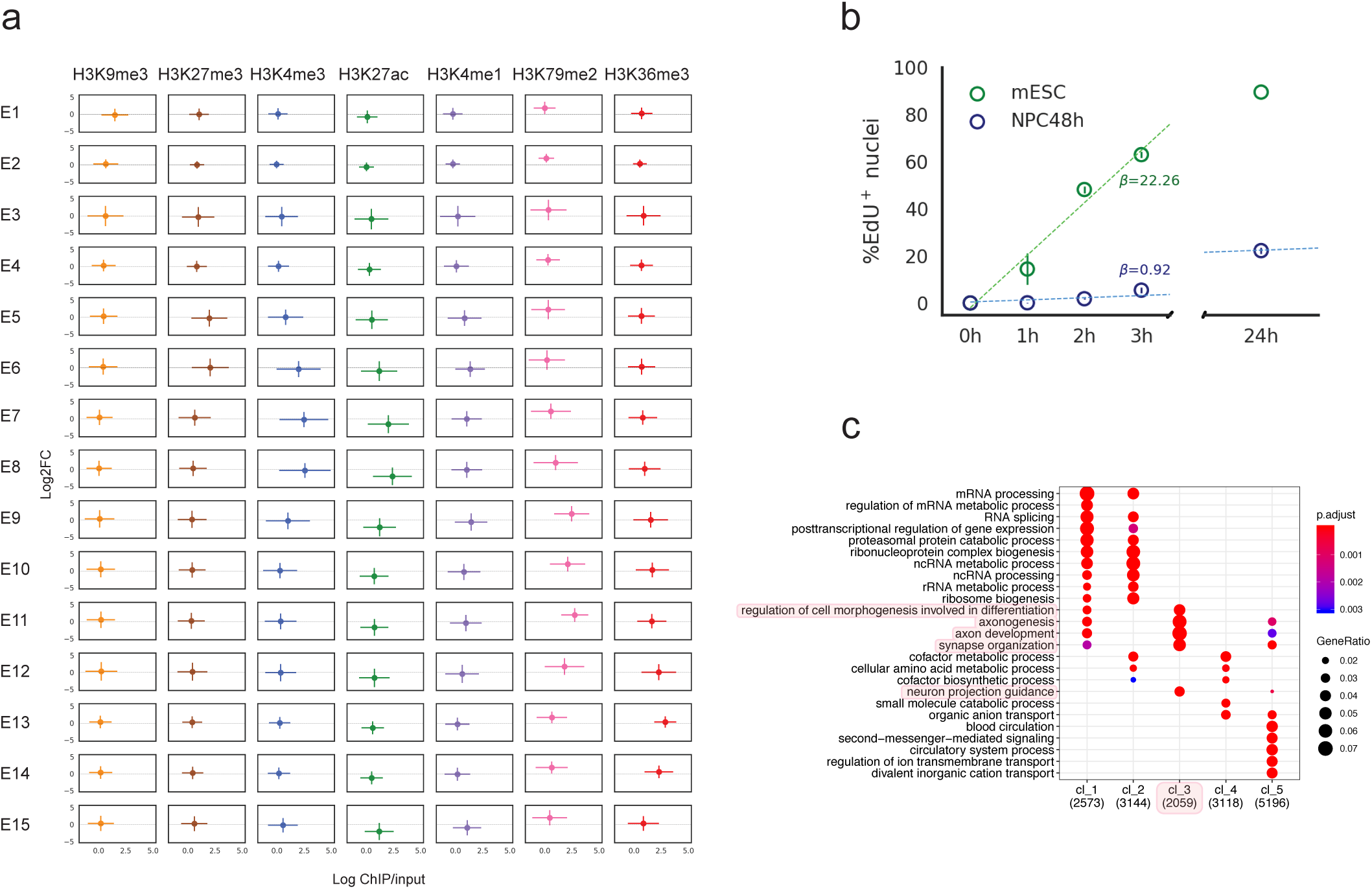
Global differences in H3K79me2 may be explained by differences in proliferation rate while local increase of the mark is detected on genes critical for neuronal differentiation. **a)** This plot generalizes the gene-centric analysis of Figure 2c. Locus-specific estimation of global log2 fold-changes of each histone modification over each mESC chromatin state segment (E1 to E15), genome-wide. H3K27ac and H3K79me3 are the only marks to show consistent changes across all chromatin states. **b)** Proliferation assay of mESC and NPC48h after EdU pulse labeling and subsequent FACS-based quantification. Percentage of EdU positive nuclei over the total number of intact nuclei (y-axis) after 1h, 2h, 3h and 24h incubation (x-axis), for mESC (green) and NPC48h (blue), n=3. **c)** Gene ontology term enrichment analysis for each of the five clusters of genes of Figure 2d. The size of each dot is proportional to the gene ratio for each significant GO term, while the color maps the adjusted p-value for the over-representation test.

**Supplementary 3.**
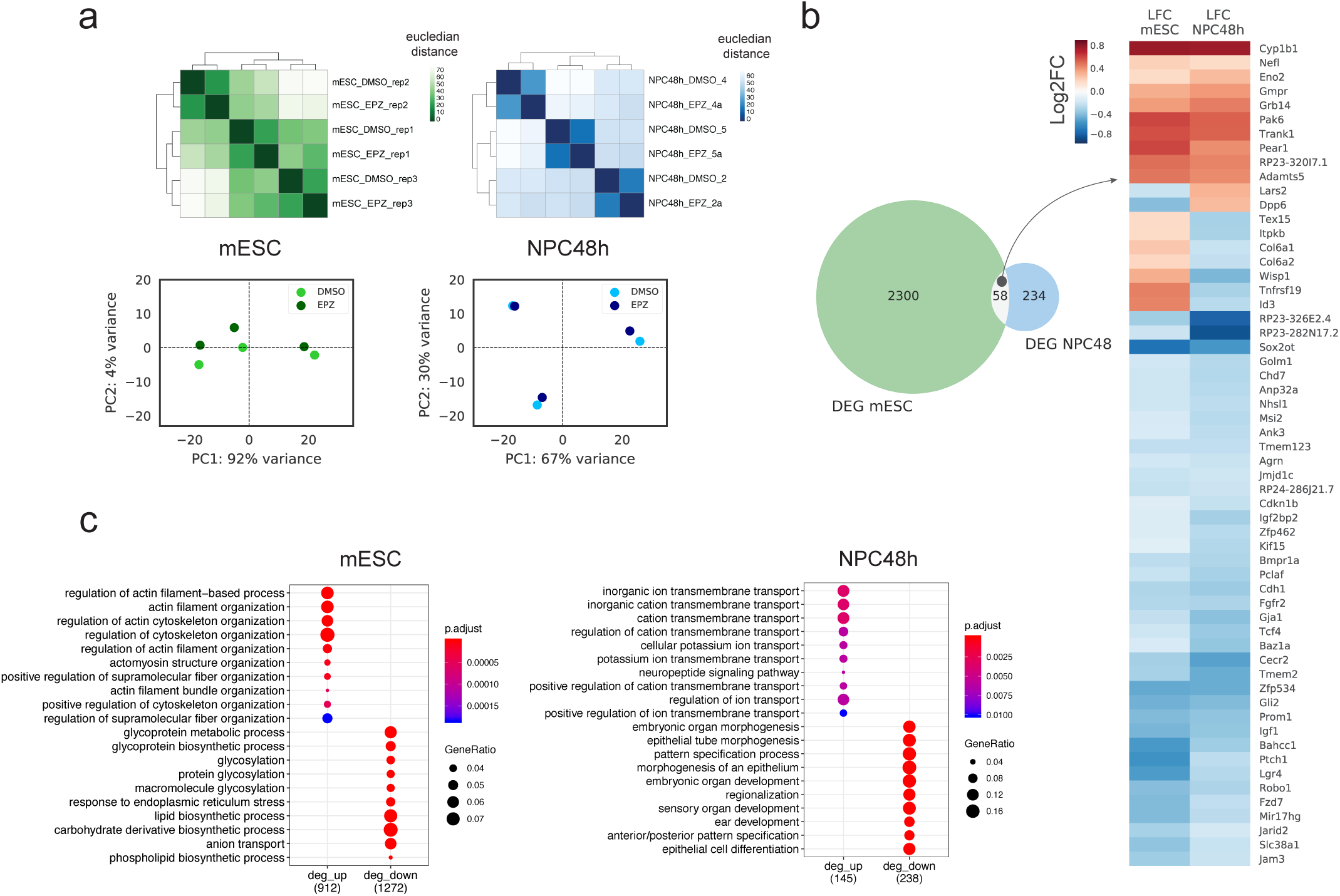
DOT1L inhibition induces cell-type specific transcriptional changes. **a)** Top panel: hierarchical clustering of mESC (green, left) and NPC48h (blue, right) samples based on the euclidean distance of the rlog transformed transcriptome. Darker shades are used for closer samples. Lower panel: principal component analysis of mESC (left, green) and NPC48h (right, blue) samples based on the top 500 most variable genes (rlog transformed counts). **b)** Left panel: intersect between EPZ-induced differential expression in mESC and NPC48h (Venn diagram on the left). Right panel: heatmap showing the log2 fold-change in expression of the 58 common differentially expressed genes in mESC and NPC48h upon DOT1L inhibition. **c)** Gene ontology term enrichment analysis for differentially expressed genes in mESC (left panel) and NPC48h (right panel) induced by EPZ treatment with respect to DMSO control. The number of genes of the upregulated and downregulated set contributing to the test is shown in parenthesis. The size of each dot is proportional to the gene ratio for each significant GO term, while the color maps the adjusted p-value for the over-representation test. deg_up: differentially upregulated genes; deg_down: differentially downregulated genes.

**Supplementary 4.**
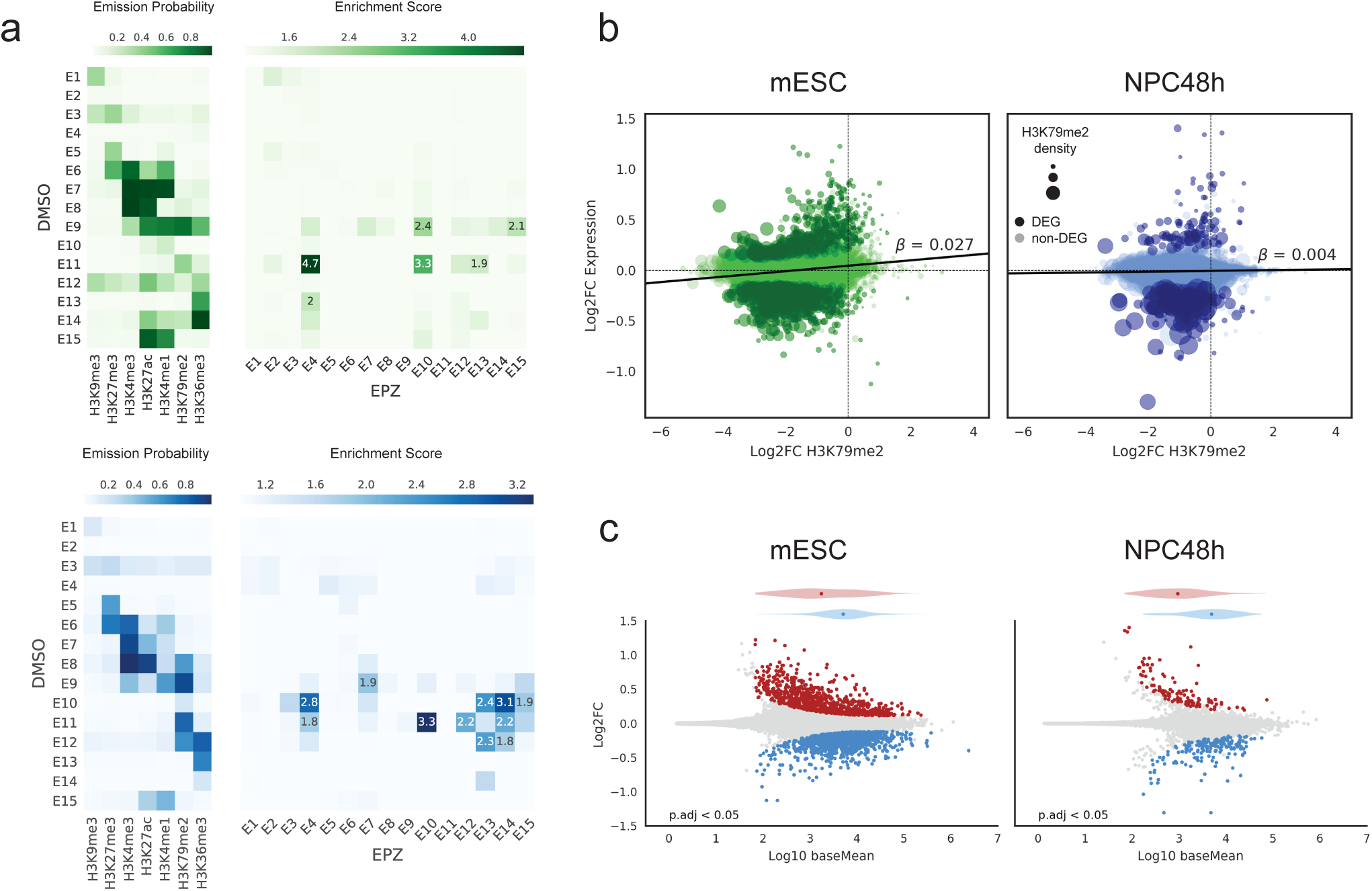
Inhibition of DOT1L does not affect the overall epigenome and transcriptome. **a)** Top panel: emission probability of the learned hidden markov model for mESC (left), next to a heatmap showing the enrichment for the transition of each state in the DMSO segmentation towards each state in the EPZ segmentation (enrichment score >= 1.9 is annotated). Lower panel shows the same analysis for NPC48h. **b)** Simple linear regression of differential expression induced by EPZ treatment compared to control on differential H3K79me2 (estimation based on RELACS data on a 3kb window downstream of TSS), for mESC (left panel, green) and NPC48h (right panel, blue). The size of each dot is proportional to the H3K79me2 density. A darker shade is used for differentially expressed genes (DEG) upon DOT1L inactivation. **c)** MA plot showing log10 of the mean count (x-axis) vs expression log2 fold-change (y-axis) as computed by DESeq2, for each gene for the contrast EPZ vs DMSO treated cells. Genes significantly increasing and decreasing in expression (adjusted p-value < 0.05) are shown in red and blue respectively. Horizontal violin plots show the Log10 mean count distribution of significantly upregulated and downregulated genes. Log2 fold-change are shrunken using apeglm. MA plot of mESC is shown on the left, while MA plot of NPC48h is shown on the right. Above each MA plot,

**Supplementary 5.**
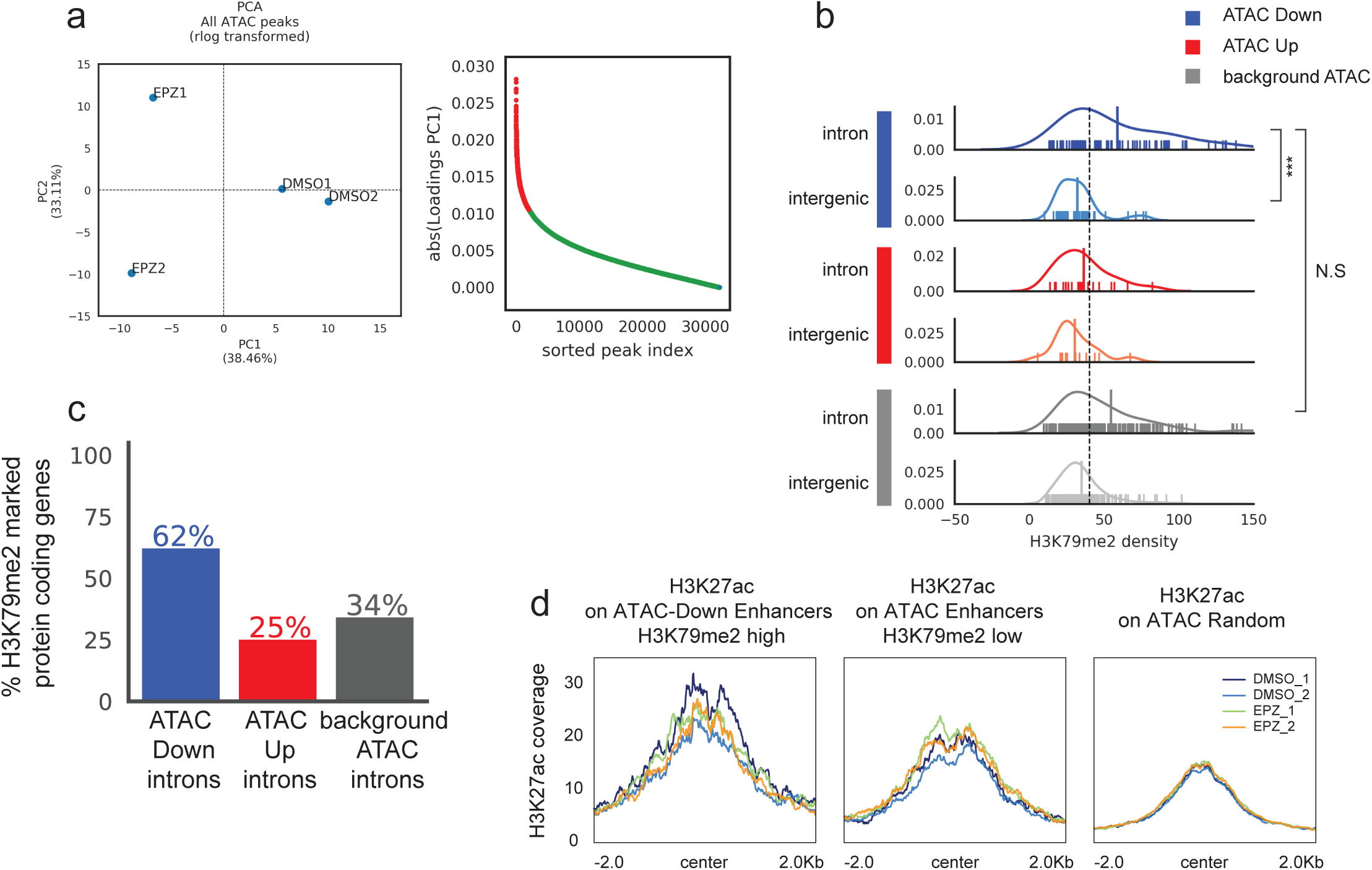
Presence of H3K79me2 is not predictive for decreased accessibility upon EPZ treatment and altered H3K79me2 do not correlate with altered H3K27ac on enhancer regions. **a)** Principal component analysis of rlog transformed ATAC peaks coverage (left panel) and ranking of ATAC peaks based on the absolute value of their PC1 loadings (right panel). The 2000 most dynamic peaks that were selected for further analysis are shown as red dots. **b)** Rug plot showing the distribution of H3K79me2 density on intergenic and intronic ATAC peaks with decreased accessibility upon EPZ treatment (ATAC-Down, blue), with increased accessibility upon EPZ treatment (ATAC-Up, red), and with no effect by EPZ treatment (background-ATAC, grey). The mean of each group is represented by the highest spike. To test for significant differences in the mean, we used the non-parametric Mann-Whitney U-test. N.S : not significant; *** : p-value < 0.001. **c)** Barplot showing the percentage of H3K79me2 positive protein coding genes over the total number of protein coding genes annotated with at least one intronic ATAC peak showing decreased accessibility (ATAC Down introns), increased accessibility (ATAC Up introns) and non-dynamic accessibility (background-ATAC introns) upon EPZ treatment with respect to DMSO control in NPC48h. **d)** Metaprofiles of H3K27ac over enhancers regions, stratified on H3K79me2 density. From left to right, we show H3K27ac profile on ATAC Down regions with high H3K79me2 (H3K79me2 density > 45), on ATAC enhancers with low H3K79me2 (H3K79me2 density < 45), and on random ATAC regions as background. Regions are aligned on the ATAC peak center.

## REFERENCES

1. Jenuwein T, Allis CD. Translating the histone code. Science. 2001 Aug 10;293(5532):1074–80.

2. Atlasi Y, Stunnenberg HG. The interplay of epigenetic marks during stem cell differentiation and development. Nat Rev Genet. 2017 Nov;18(11):643–58.

3. Juliandi B, Abematsu M, Nakashima K. Epigenetic regulation in neural stem cell differentiation. Dev Growth Differ. 2010 Aug;52(6):493–504.

4. Podobinska M, Szablowska-Gadomska I, Augustyniak J, Sandvig I, Sandvig A, Buzanska L. Epigenetic Modulation of Stem Cells in Neurodevelopment: The Role of Methylation and Acetylation. Front Cell Neurosci. 2017 Feb 7;11:23.

5. Efroni S, Duttagupta R, Cheng J, Dehghani H, Hoeppner DJ, Dash C, et al. Global transcription in pluripotent embryonic stem cells. Cell Stem Cell. 2008 May 8;2(5):437–47.

6. Arrigoni L, Al-Hasani H, Ramírez F, Panzeri I, Ryan DP, Santacruz D, et al. RELACS nuclei barcoding enables high-throughput ChIP-seq. Commun Biol. 2018 Dec 5;1:214.

7. Orlando DA, Chen MW, Brown VE, Solanki S, Choi YJ, Olson ER, et al. Quantitative ChIP-Seq normalization reveals global modulation of the epigenome. Cell Rep. 2014 Nov 6;9(3):1163–70.

8. van Galen P, Viny AD, Ram O, Ryan RJH, Cotton MJ, Donohue L, et al. A Multiplexed System for Quantitative Comparisons of Chromatin Landscapes. Mol Cell. 2016 Jan 7;61(1):170–80.

9. Boland MJ, Nazor KL, Loring JF. Epigenetic regulation of pluripotency and differentiation. Circ Res. 2014 Jul 7;115(2):311–24.

10. Onder TT, Kara N, Cherry A, Sinha AU, Zhu N, Bernt KM, et al. Chromatin-modifying enzymes as modulators of reprogramming. Nature. 2012 Mar 4;483(7391):598–602.

11. Pursani V, Bhartiya D, Tanavde V, Bashir M, Sampath P. Transcriptional activator DOT1L putatively regulates human embryonic stem cell differentiation into the cardiac lineage. Stem Cell Res Ther. 2018 Apr 10;9(1):97.

12. Barry ER, Krueger W, Jakuba CM, Veilleux E, Ambrosi DJ, Nelson CE, et al. ES cell cycle progression and differentiation require the action of the histone methyltransferase Dot1L. Stem Cells. 2009 Jul;27(7):1538–47.

13. Franz H, Villarreal A, Heidrich S, Videm P, Kilpert F, Mestres I, et al. DOT1L promotes progenitor proliferation and primes neuronal layer identity in the developing cerebral cortex. Nucleic Acids Res. 2019 Jan 10;47(1):168–83.

14. Wood K, Tellier M, Murphy S. DOT1L and H3K79 Methylation in Transcription and Genomic Stability. Biomolecules [Internet]. 2018 Feb 27;8(1). Available from: http://dx.doi.org/10.3390/biom8010011

15. van Leeuwen F, Gafken PR, Gottschling DE. Dot1p modulates silencing in yeast by methylation of the nucleosome core. Cell. 2002 Jun 14;109(6):745–56.

16. Singer MS, Kahana A, Wolf AJ, Meisinger LL, Peterson SE, Goggin C, et al. Identification of high-copy disruptors of telomeric silencing in Saccharomyces cerevisiae. Genetics. 1998 Oct;150(2):613–32.

17. Kim W, Choi M, Kim J-E. The histone methyltransferase Dot1/DOT1L as a critical regulator of the cell cycle. Cell Cycle. 2014 Feb 6;13(5):726–38.

18. Steger DJ, Lefterova MI, Ying L, Stonestrom AJ, Schupp M, Zhuo D, et al. DOT1L/KMT4 recruitment and H3K79 methylation are ubiquitously coupled with gene transcription in mammalian cells. Mol Cell Biol. 2008 Apr;28(8):2825–39.

19. McLean CM, Karemaker ID, van Leeuwen F. The emerging roles of DOT1L in leukemia and normal development. Leukemia. 2014 Nov;28(11):2131–8.

20. Cattaneo P, Kunderfranco P, Greco C, Guffanti A, Stirparo GG, Rusconi F, et al. DOT1L-mediated H3K79me2 modification critically regulates gene expression during cardiomyocyte differentiation. Cell Death Differ. 2016 Apr;23(4):555–64.

21. Nguyen AT, Xiao B, Neppl RL, Kallin EM, Li J, Chen T, et al. DOT1L regulates dystrophin expression and is critical for cardiac function. Genes Dev. 2011 Feb 1;25(3):263–74.

22. Castaño Betancourt MC, Cailotto F, Kerkhof HJ, Cornelis FMF, Doherty SA, Hart DJ, et al. Genome-wide association and functional studies identify the DOT1L gene to be involved in cartilage thickness and hip osteoarthritis. Proc Natl Acad Sci U S A. 2012 May 22;109(21):8218–23.

23. Roidl D, Hellbach N, Bovio PP, Villarreal A, Heidrich S, Nestel S, et al. DOT1L Activity Promotes Proliferation and Protects Cortical Neural Stem Cells from Activation of ATF4-DDIT3-Mediated ER Stress In Vitro. Stem Cells. 2016 Jan;34(1):233–45.

24. Bovio PP, Franz H, Heidrich S, Rauleac T, Kilpert F, Manke T, et al. Differential Methylation of H3K79 Reveals DOT1L Target Genes and Function in the Cerebellum In Vivo. Mol Neurobiol. 2019 Jun;56(6):4273–87.

25. Chen C-W, Koche RP, Sinha AU, Deshpande AJ, Zhu N, Eng R, et al. DOT1L inhibits SIRT1-mediated epigenetic silencing to maintain leukemic gene expression in MLL-rearranged leukemia. Nat Med. 2015 Apr;21(4):335–43.

26. Godfrey L, Crump NT, Thorne R, Lau I-J, Repapi E, Dimou D, et al. DOT1L inhibition reveals a distinct subset of enhancers dependent on H3K79 methylation. Nat Commun. 2019 Jun 26;10(1):2803.

27. Bibel M, Richter J, Lacroix E, Barde Y-A. Generation of a defined and uniform population of CNS progenitors and neurons from mouse embryonic stem cells. Nat Protoc. 2007;2(5):1034–43.

28. Cheng C, Gerstein M. Modeling the relative relationship of transcription factor binding and histone modifications to gene expression levels in mouse embryonic stem cells. Nucleic Acids Res. 2012 Jan;40(2):553–68.

29. Montavon T, Soshnikova N. Hox gene regulation and timing in embryogenesis. Semin Cell Dev Biol. 2014 Oct;34:76–84.

30. El Wazan L, Urrutia-Cabrera D, Wong RC-B. Using transcription factors for direct reprogramming of neurons in vitro. World J Stem Cells. 2019 Jul 26;11(7):431–44.

31. Ernst J, Kellis M. ChromHMM: automating chromatin-state discovery and characterization. Nat Methods. 2012 Feb 28;9(3):215–6.

32. Ernst J, Kellis M. Chromatin-state discovery and genome annotation with ChromHMM. Nat Protoc. 2017 Dec;12(12):2478–92.

33. De Vos D, Frederiks F, Terweij M, van Welsem T, Verzijlbergen KF, Iachina E, et al. Progressive methylation of ageing histones by Dot1 functions as a timer. EMBO Rep. 2011 Sep 1;12(9):956–62.

34. ENCODE Project Consortium. An integrated encyclopedia of DNA elements in the human genome. Nature. 2012 Sep 6;489(7414):57–74.

35. Chory EJ, Calarco JP, Hathaway NA, Bell O, Neel DS, Crabtree GR. Nucleosome Turnover Regulates Histone Methylation Patterns over the Genome. Mol Cell. 2019 Jan 3;73(1):61– 72.e3.

36. Zywitza V, Misios A, Bunatyan L, Willnow TE, Rajewsky N. Single-Cell Transcriptomics Characterizes Cell Types in the Subventricular Zone and Uncovers Molecular Defects Impairing Adult Neurogenesis. Cell Rep. 2018 Nov 27;25(9):2457–69.e8.

37. Loo L, Simon JM, Xing L, McCoy ES, Niehaus JK, Guo J, et al. Single-cell transcriptomic analysis of mouse neocortical development. Nat Commun. 2019 Jan 11;10(1):134.

38. Lu TT-H, Heyne S, Dror E, Casas E, Leonhardt L, Boenke T, et al. The Polycomb-Dependent Epigenome Controls β Cell Dysfunction, Dedifferentiation, and Diabetes. Cell Metab. 2018 Jun 5;27(6):1294–308.e7.

39. Dennis DJ, Han S, Schuurmans C. bHLH transcription factors in neural development, disease, and reprogramming. Brain Res. 2019 Feb 15;1705:48–65.

40. Park NI, Guilhamon P, Desai K, McAdam RF, Langille E, O’Connor M, et al. ASCL1 Reorganizes Chromatin to Direct Neuronal Fate and Suppress Tumorigenicity of Glioblastoma Stem Cells. Cell Stem Cell. 2017 Aug 3;21(2):209–24.e7.

41. Heinz S, Romanoski CE, Benner C, Glass CK. The selection and function of cell type-specific enhancers. Nat Rev Mol Cell Biol. 2015 Mar;16(3):144–54.

42. Thiel G. How Sox2 maintains neural stem cell identity. Biochem J. 2013 Mar 15;450(3):e1– 2.

43. Pevny LH, Nicolis SK. Sox2 roles in neural stem cells. Int J Biochem Cell Biol. 2010 Mar;42(3):421–4.

44. Chen C, Lee GA, Pourmorady A, Sock E, Donoghue MJ. Orchestration of Neuronal Differentiation and Progenitor Pool Expansion in the Developing Cortex by SoxC Genes. J Neurosci. 2015 Jul 22;35(29):10629–42.

45. Bertolini JA, Favaro R, Zhu Y, Pagin M, Ngan CY, Wong CH, et al. Mapping the Global Chromatin Connectivity Network for Sox2 Function in Neural Stem Cell Maintenance. Cell Stem Cell. 2019 Mar 7;24(3):462–76.e6.

46. Ugarte F, Sousae R, Cinquin B, Martin EW, Krietsch J, Sanchez G, et al. Progressive Chromatin Condensation and H3K9 Methylation Regulate the Differentiation of Embryonic and Hematopoietic Stem Cells. Stem Cell Reports. 2015 Nov 10;5(5):728–40.

47. Chen T, Dent SYR. Chromatin modifiers and remodellers: regulators of cellular differentiation. Nat Rev Genet. 2014 Feb;15(2):93–106.

48. Meshorer E, Yellajoshula D, George E, Scambler PJ, Brown DT, Misteli T. Hyperdynamic plasticity of chromatin proteins in pluripotent embryonic stem cells. Dev Cell. 2006 Jan;10(1):105–16.

49. Mattout A, Meshorer E. Chromatin plasticity and genome organization in pluripotent embryonic stem cells. Curr Opin Cell Biol. 2010 Jun;22(3):334–41.

50. Bennett CG, Riemondy K, Chapnick DA, Bunker E, Liu X, Kuersten S, et al. Genome-wide analysis of Musashi-2 targets reveals novel functions in governing epithelial cell migration. Nucleic Acids Res. 2016 May 5;44(8):3788–800.

51. Li G, Margueron R, Ku M, Chambon P, Bernstein BE, Reinberg D. Jarid2 and PRC2, partners in regulating gene expression. Genes Dev. 2010 Feb 15;24(4):368–80.

52. Pasini D, Cloos PAC, Walfridsson J, Olsson L, Bukowski J-P, Johansen JV, et al. JARID2 regulates binding of the Polycomb repressive complex 2 to target genes in ES cells. Nature. 2010 Mar 11;464(7286):306–10.

53. Kwak S, Kim TW, Kang B-H, Kim J-H, Lee J-S, Lee H-T, et al. Zinc finger proteins orchestrate active gene silencing during embryonic stem cell differentiation. Nucleic Acids Res. 2018 Jul 27;46(13):6592–607.

54. Arrigoni L, Richter AS, Betancourt E, Bruder K, Diehl S, Manke T, et al. Standardizing chromatin research: a simple and universal method for ChIP-seq. Nucleic Acids Res. 2016 Apr 20;44(7):e67.

55. Buenrostro JD, Wu B, Chang HY, Greenleaf WJ. ATAC-seq: A Method for Assaying Chromatin Accessibility Genome-Wide. Curr Protoc Mol Biol. 2015 Jan 5;109:21.29.1–9.

56. Abramoff, Magalhães PJ, Ram SJ. Image processing with ImageJ. Biophotonics international. 2004;11(7):36–42.

57. Bhardwaj V, Heyne S, Sikora K, Rabbani L, Rauer M, Kilpert F, et al. snakePipes enable flexible, scalable and integrative epigenomic analysis [Internet]. bioRxiv. 2018 [cited 2019 Apr 11]. p. 407312. Available from: https://www.biorxiv.org/content/10.1101/407312v2

58. Smith T, Heger A, Sudbery I. UMI-tools: modeling sequencing errors in Unique Molecular Identifiers to improve quantification accuracy. Genome Res. 2017 Mar;27(3):491–9.

59. Love MI, Huber W, Anders S. Moderated estimation of fold change and dispersion for RNA-seq data with DESeq2. Genome Biol. 2014;15(12):550.

60. Zhu A, Ibrahim JG, Love MI. Heavy-tailed prior distributions for sequence count data: removing the noise and preserving large differences [Internet]. bioRxiv. 2018 [cited 2019 Apr 11]. p. 303255. Available from: https://www.biorxiv.org/content/10.1101/303255v1

61. Ramírez F, Ryan DP, Grüning B, Bhardwaj V, Kilpert F, Richter AS, et al. deepTools2: a next generation web server for deep-sequencing data analysis. Nucleic Acids Res. 2016 Jul 8;44(W1):W160–5.

62. Heinz S, Benner C, Spann N, Bertolino E, Lin YC, Laslo P, et al. Simple combinations of lineage-determining transcription factors prime cis-regulatory elements required for macrophage and B cell identities. Mol Cell. 2010 May 28;38(4):576–89.

63. Kondili M, Fust A, Preussner J, Kuenne C, Braun T, Looso M. UROPA: a tool for Universal RObust Peak Annotation. Sci Rep. 2017 Jun 1;7(1):2593.

64. van Heeringen SJ, Veenstra GJC. GimmeMotifs: a de novo motif prediction pipeline for ChIP-sequencing experiments. Bioinformatics. 2011 Jan 15;27(2):270–1.

65. Richard G. gtrichard/deepStats: deepStats 0.3.1 [Internet]. 2019. Available from: https://zenodo.org/record/3361799

66. Lê S, Josse J, Husson F. FactoMineR: An R package for multivariate analysis. 2008 [cited 2019 Nov 5]; Available from: https://hal.archives-ouvertes.fr/hal-00359835

67. Yu G, Wang L-G, Han Y, He Q-Y. clusterProfiler: an R package for comparing biological themes among gene clusters. OMICS. 2012 May;16(5):284–7.

68. Maaten L van der, Hinton G. Visualizing Data using t-SNE. J Mach Learn Res. 2008;9(Nov):2579–605.

69. Salvatier J, Wiecki TV, Fonnesbeck C. Probabilistic programming in Python using PyMC3. PeerJ Comput Sci. 2016 Apr 6;2:e55.

